# Seafloor incubation experiment with deep-sea hydrothermal vent fluid reveals effect of pressure and lag time on autotrophic microbial communities

**DOI:** 10.1101/2020.11.11.378315

**Authors:** Caroline S. Fortunato, David A. Butterfield, Benjamin Larson, Noah Lawrence-Slavas, Christopher K. Algar, Lisa Zeigler Allen, James F. Holden, Giora Proskurowski, Emily Reddington, Lucy C. Stewart, Begüm D. Topçuoğlu, Joseph J. Vallino, Julie A. Huber

**Author notes:** To whom correspondence should be directed.

## Abstract

Depressurization and sample processing delays may impact the outcome of shipboard microbial incubations of samples collected from the deep sea. To address this knowledge gap, we developed an ROV-powered incubator instrument to carry out and compare results from *in situ* and shipboard RNA Stable Isotope Probing (RNA-SIP) experiments to identify the key chemolithoautotrophic microbes and metabolisms in diffuse, low-temperature venting fluids from Axial Seamount. All the incubations showed microbial uptake of labelled bicarbonate primarily by thermophilic autotrophic Epsilonbacteraeota that oxidized hydrogen coupled with nitrate reduction. However, the *in situ* seafloor incubations showed higher abundances of transcripts annotated for aerobic processes suggesting that oxygen was lost from the hydrothermal fluid samples prior to shipboard analysis. Furthermore, transcripts for thermal stress proteins such as heat shock chaperones and proteases were significantly more abundant in the shipboard incubations suggesting that hydrostatic pressure ameliorated thermal stress in the metabolically active microbes in the seafloor incubations. Together, results indicate that while the autotrophic microbial communities in the shipboard and seafloor experiments behaved similarly, there were distinct differences that provide new insight into the activities of natural microbial assemblages under near-native conditions in the ocean.

## Introduction

At deep-sea hydrothermal vents, low temperature (i.e., diffuse) hydrothermal fluids emanating directly from igneous rock are hot spots of microbial primary production and provide access points to subseafloor habitats. Diffuse vents are formed when cold, oxidized seawater mixes with hot chemically reduced hydrothermal fluids at and below the seafloor, creating steep geochemical gradients that support increased microbial biomass, activity, and diversity relative to the surrounding deep ocean (Butterfield et al 2004, Huber et al 2007, Jannasch and Mottl 1985, McNichol et al 2018, Perner et al 2009). These fluids are dominated by chemolithoautotrophic bacteria and archaea that carry out a variety of metabolisms utilizing hydrogen, sulfur compounds, nitrate, and methane (Fortunato and Huber 2016, Fortunato et al 2018, Galambos et al 2019, Meier et al 2017, Olins et al 2017, Reveillaud et al 2016, Trembath-Reichert et al 2019). However, our understanding of the impact of different microbial metabolisms on ocean biogeochemistry and the extent of carbon production from these reactions are nascent. This is partially due to the challenges associated with the collection and transfer of samples from the deep ocean to the surface for experimentation.

Samples transferred from the deep ocean to the sea surface are subject to changes in temperature and pressure and usually involve a long lag time between collection, sample recovery, and shipboard processing. Deep-sea devices designed for filtering seawater and other fluids at depth have been used to minimize these issues through *in situ* filtration, cell concentration, preservation, and analysis (reviewed in Edgcomb et al 2016, Ottesen 2016). The outgassing of compounds such as hydrogen and carbon dioxide impacts microbial measurements from deep-sea hydrothermal vents; therefore, samples often need to be maintained at *in situ* pressures or temperatures when possible (reviewed in Sievert and Vetriani 2012). For example, McNichol et al. (2016, 2018) used an isobaric gas tight fluid sampler to conduct shipboard carbon fixation experiments with diffuse vent fluids maintained at *in situ* pressures and temperatures. While such measurements remain critical to constraining microbial processes in the ocean, these and other such experiments have sample processing delays and lack *in situ* preservation. This could be critical when sampling microbial communities in diffuse fluids that are in an extreme state of chemical disequilibrium and will likely undergo redox reactions between sampling and arrival in shipboard labs, regardless of the temperature and pressure conditions maintained in the sampling device.

A limited suite of samplers have been developed to carry out experiments while deployed in the ocean, keeping the instrument submerged for the duration of the experiment and fixing the samples post-experiment, before instrument recovery. This avoids biases related to sample collection lag and depressurization, although other experimental artifacts, such as bottle effects still remain (reviewed in McQuillan and Robidart 2017, Ottesen 2016). This includes the automated micro-laboratory designed to allow one to conduct multiple (in-series) tracer incubation studies during cabled or free-drifting deployments (Lippsett 2014, Taylor et al 1993, Taylor et al 1983, Taylor and Doherty 1990), as well as a modification of the instrument termed the Microbial Sampler-Submersible Incubation Device (MS-SID), allowing for in situ grazing incubation experiments together with in situ microbial sampling and preservation (Pachiadaki et al 2016; Edgcomb et al 2016; Medina et al 2017). Another instrument is the Environmental Sample Processor (ESP) unit which includes a molecular component that carries out sample homogenization and subsequent detection of particular microbial groups using quantitative PCR, sandwich hybridization, or competitive ELISA (Scholin 2018). A version of the ESP has successfully been deployed in the deep ocean, including in venting hydrothermal fluids (Olins et al 2017) and methane seeps (Ussler et al 2013).

We recently developed a shipboard RNA Stable Isotope Probing (RNA-SIP) procedure combined with metatranscriptomics to identify the key chemolithoautotrophs and metabolisms present in deep-sea hydrothermal vent ecosystems (Fortunato and Huber 2016, Trembath-Reichert et al 2019). In this study, the method was extended to the seafloor by running RNA-SIP experiments in a newly developed incubator that collects, heats, incubates, manipulates, and preserves seawater and vent fluids to allow for *in situ* experimentation while powered by a remotely operated vehicle. The results of the *in situ* incubation experiment were compared with parallel shipboard experiments in order to determine the effect of pressure changes and lag time on microbial metabolism. Herein, we describe the new *in situ* incubator and the results of metatranscriptomic sequencing of the shipboard and seafloor RNA-SIP experiments to provide new insights into the activities of natural microbial assemblages under near-native conditions in the deep ocean.

## Methods

### Fluid collection

Low temperature (41°C) hydrothermal vent fluid was collected from Marker 33 vent at Axial Seamount (45.93346, −129.98225, 1516 m depth) on 26 August 2015 on board the *R/V Thomas G. Thompson* using *ROV Jason* II. Fluids were collected using the Hydrothermal Fluid and Particle Sampler (HFPS, Butterfield et al 2004), which has an integrated temperature sensor to continuously monitor fluid temperature during intake. Collection and processing of diffuse vent fluid samples for RNA-SIP are described below. For collection of filtered vent fluid for microbial community DNA and RNA analyses, 3 L of diffuse fluid was pumped through a 0.22 μm pore size, 47 mm diameter GWSP filter and preserved immediately *in situ* with RNALater as described previously (Fortunato et al. 2018). Separate fluid samples were collected and analyzed for alkalinity and hydrogen sulfide, ammonia, methane, and hydrogen concentrations following methods described previously (Butterfield et al. 2004). The oxygen concentration and pH of the fluid were measured during intake using a Seabird 63 Optical oxygen sensor and an AMT deep-sea glass pH electrode that were integrated into the HFPS.

### Shipboard RNA stable isotope probing experiments

Shipboard RNA-SIP experiments were performed as previously described (Fortunato and Huber 2016). The HFPS was used to collect 4 L of diffuse vent fluid into an acid washed Tedlar bag. Once on the ship, diffuse fluid was pumped from the Tedlar bag into four evacuated 500 mL Pyrex bottles and filled to capacity (530 mL). Prior to filling, ^12^C-labeled sodium bicarbonate or ^13^C sodium bicarbonate was added separately to a pair of bottles to reach a final added concentration of 10 mM bicarbonate. After adding the fluid sample to each bottle, 1 mL of 1.2 M HCl was added to counteract the added bicarbonate and ensure a pH similar to unamended vent fluid. H_2_ (900 μmol) was then added to each bottle. A pair of ^13^C- and ^12^C-labelled bottles was then incubated at 55°C for 12 h while another pair was incubated for 16 h. After incubation, the fluid from each bottle was filtered separately through 0.22 μm pore size Sterivex filters, preserved in RNALater, and frozen at −80°C.

### Seafloor RNA stable isotope probing experiments

The seafloor incubator units were incorporated as a module on the HFPS (Figure 1) and designed to pull in vent fluid using the existing HFPS framework and was A/C powered by the submersible. The incubations occurred concurrently with other HFPS fluid collection and dive operations. The main components of a single incubator unit consisted of an insulated incubator bottle containing the primary sample bag (4 mil thick Tedlar bag), an RTD (Resistance Temperature Detector) probe, and a 250 W heating rod and a final bottle containing a secondary sample bag (2 mil thick Tedlar bag) and a titanium shutoff valve situated between the fluid intake lines and incubator bottle. Four insulated incubation units were loaded onto one rack of the HFPS (Figure 1).

**Figure 1.**
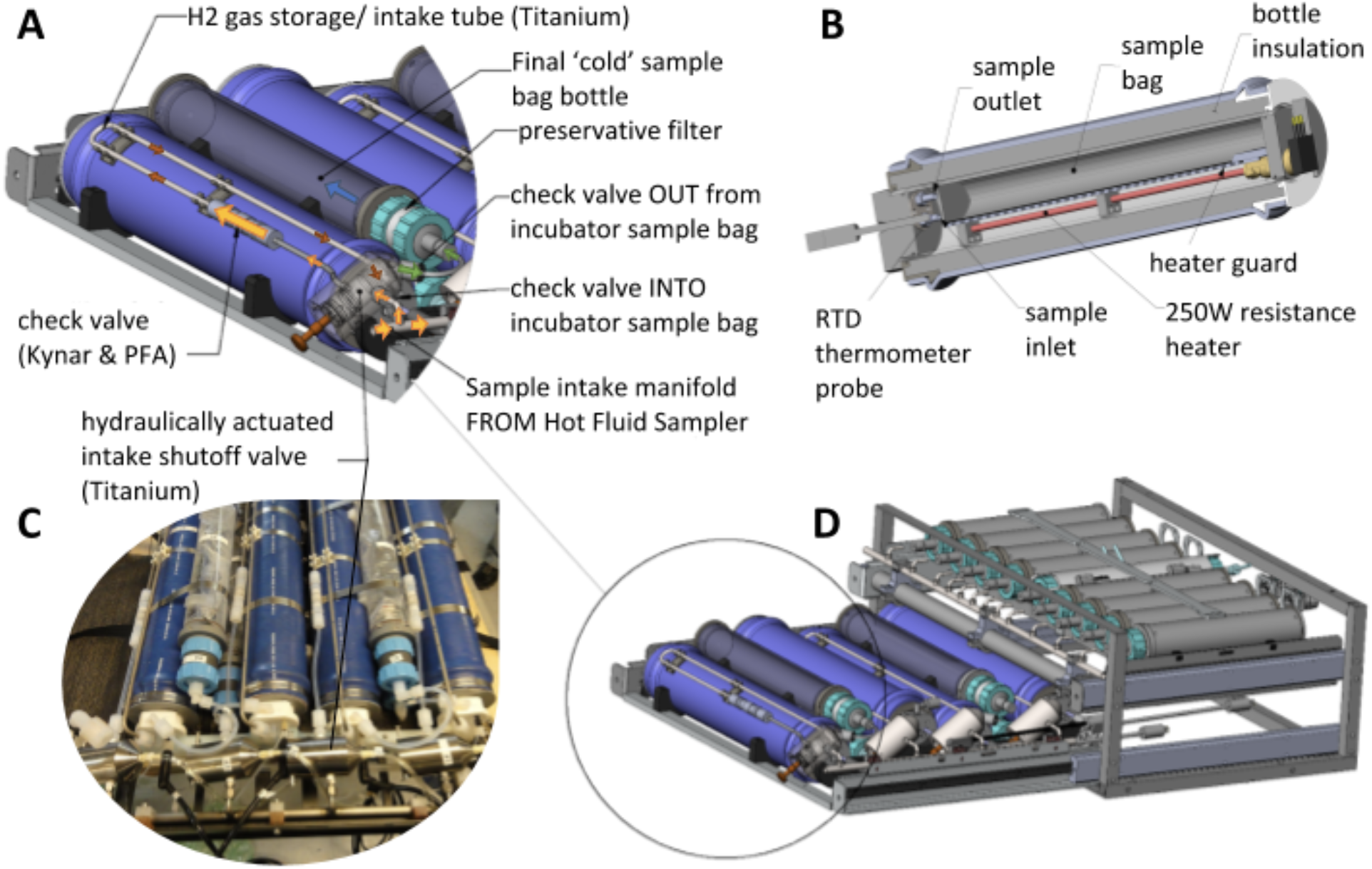
Incubator setup for the *in situ* RNA Stable Isotope Probing (RNA-SIP) experiments (A-D). Each of the four incubation chambers was heated to a chosen set point temperature. Fluid was pulled into the insulated incubation chamber from the manifold of the Hydrothermal Fluid and Particle Sampler (HFPS) through a custom titanium shutoff valve, pulling hydrogen gas and buffering acid into the chamber as it filled. After the incubation period, the fluid was pulled from the incubation chamber through a 0.22 μm filter (A) with passive addition of RNA preservative. A cutaway view of the incubation chamber (B) shows the incubation bag over the heating element, with the RTD used to monitor chamber temperature near the end of the bag. The fully assembled incubator module (C, as deployed in 2015) slides into the HFPS sample rack (D). Fluid transfer is accomplished with the HFPS sample pump and selection valve.

Prior to deployment, ^12^C-labeled sodium bicarbonate or ^13^C sodium bicarbonate was added separately to a pair each of primary incubation bags to reach a final concentration of 10 mM added bicarbonate upon filling with 800 mL of vent fluid. The lines running to each bag were primed with 1.5 mL of 1.2 M HCl to ensure a pH similar to unamended vent fluid as well as 900 μmol of pure H_2_ to match the shipboard incubations. Approximately one hour prior to fluid sampling on the seafloor, the insulated incubator chambers were heated to 55°C. This incubation temperature was selected based on the high abundances of thermophiles at Marker 33 in previous studies (Fortunato et al 2018, Huber et al 2003). Once at temperature, the primary sample bags were filled with 800 mL of diffuse vent fluid using the HFPS as described above and a shut off valve was hydraulically closed to prevent further intake from the sample manifold. An RKC MA901 Proportional-Integral-Derivative (PID) temperature controller housed in a separate titanium case recorded and controlled incubator temperature from an RTD thermometer situated next to the bag and maintained a constant temperature at a set point (± 2°C) by supplying variable power to the heating rod located beneath the Tedlar incubation bag inside the incubator (Figure S1). The PID control algorithm was tuned to the incubator bottle prior to deployment using the MA901 autotune feature. The heating rod induced convection in the incubation chamber resulting in an even temperature distribution. The temperature distribution within the incubator sample bag was monitored during pre-deployment laboratory experiments and was found to vary less than 2°C (Table S3).

A pair of insulated incubator chambers containing ^12^C and ^13^C bicarbonate were incubated for 12 h while an identical pair of chambers were incubated for 16 h. At the end of each incubation, fluid was pumped from the primary incubator bag, through a 0.22 μm pore size PES filter (Millipore) into a secondary bag that was surrounded by ambient seawater (~ 2°C). Filters were preserved immediately *in situ* with RNALater. Once shipboard, the fluids in the secondary sample bags were analyzed for pH and the filters were frozen at −80°C.

### Fractionation of RNA-SIP experiments, RT-qPCR, and library preparation

RNA from the incubator and shipboard SIP experiments was extracted, quantified, and fractionated after isopycnic centrifugation as described in Fortunato and Huber (2016) and Supplemental Material. 16S rRNA copy number was determined for each fraction via RT-qPCR with universal primers Pro341F and Pro805R (Takahashi et al 2014) as described in the Supplemental Material. This measurement was used for comparison between the ^12^C and ^13^C samples and for determination of ^13^C enrichment. Four fractions from each of the ^12^C and ^13^C samples from the shipboard and incubator samples were sequenced, including fractions with the maximum amount of 16S rRNA and fractions on either side of the peak, for a total of 16 metatranscriptomic libraries. SIP metatranscriptomic library preparation was completed as described in the Supplemental Material.

### Metagenomic and metatranscriptomic library preparation, sequencing, and analysis

The 47 mm diameter flat filters were cut in half with a sterile razor with each half used for DNA and RNA extractions, respectively, and corresponding libraries prepared for sequencing as described in the Supplemental Material and Fortunato et al. 2018. For the RNA-SIP metatranscriptomes, taxonomy, overall transcript abundance, and hierarchical clustering is displayed for all 16 libraries. For visualization of key metabolic processes, the 16 libraries were collapsed into their corresponding experiments: ^12^C Shipboard, ^13^C Shipboard, ^12^C Incubator, and ^13^C Incubator. Transcript abundance across fractions was summed for each experiment.

### Differential Expression Analysis

To determine significance between transcript abundances across RNA-SIP fractions, differential expression (DE) analysis was run using the interactive tool DEBrowser in R (Kucukural et al. 2019). Within DEBrowser, differential expression analysis was run using normalized transcript abundances for KO annotated genes for all 16 RNA-SIP libraries with Limma (Ritchie et al. 2015). Low count transcripts, defined as the maximum normalized abundance for each transcript across all samples being less than 10, were removed from the analysis. Resulting tables, heatmaps, and plots showing significance were generated within DEBrowser.

### Mapping to thermophilic Epsilonbacteraeota MAGs

Metagenome Assembled Genomes (MAGs) were assembled and taxonomically identified from Axial Seamount metagenomic data as described in Fortunato et al. (2018). In this study we determined the mean coverage of the 10 previously identified MAGs classified as thermophilic *Epsilonbacteraeota* via Phylosift (Darling et al 2014) and CheckM (Parks et al 2015) within the dataset (Table S2). The Marker 33 metagenome, Marker 33 metatranscriptome, and all RNA-SIP metatranscriptomes were mapped to each of the ten MAGs using Bowtie2 with an end-to-end alignment and default parameters (v2.0.0-beta5 Langmead and Salzberg 2012). Mean coverage for each MAG within the Marker 33 metagenome was calculated via *Anvi’o* (Eren et al 2015).

Mean coverage for each MAG within the Marker 33 metatranscriptome and RNA-SIP metatranscriptomes was calculated via *samtools* (Li et al 2009). For ease of visualization, the 16 RNA-SIP metatranscriptomes were collapsed and mean coverage for each MAG within fractions was averaged for each of the four experiments: ^12^C Shipboard, ^13^C Shipboard, ^12^C Incubator, and ^13^C Incubator. Heatmaps of mean coverage were constructed in R using the package *heatmap3* (v3.3.2, R-Development-Core-Team 2011).

### Data deposition

Raw sequence data are publicly available through the European Nucleotide Archive (ENA), with project number PRJEB38697 for RNA-SIP metatranscriptomes and PRJEB19456 for the Marker 33 metagenome and metatranscriptome. Assembled contigs for the Marker 33 metagenome and RNA-SIP metatranscriptomes are publicly available via IMG/MER under submission numbers 78401, 97537-97540, and 97583-97594. Contigs for the 10 *Epsilonbacteraeota* MAGs are available through FigShare at DOI: 10.6084/m9.figshare.12445976

## Results

### Enrichment observed in RNA-SIP experiments

Diffuse hydrothermal fluid at Marker 33 vents directly from cracks in basalt along the eruption zone on the southeast side of the Axial caldera. Chemical analysis of this fluid is shown in Table S1. The fluid was 85% seawater and 15% hydrothermal end-member fluid based on magnesium concentration (Fortunato et al 2018). The temperature was monitored throughout the experiment, and temperature records showed that the incubator rapidly heated the chambers to 55°C and maintained temperature within 2°C for the length of the incubations (Figure S1). Upon recovery, the mass of each secondary bag was determined to indicate how much incubated sample was pulled through the RNA preservative filter at the end of the incubation. In general, the secondary bags were full or nearly full and the primary incubator bags were empty or nearly empty. The pH of the filtered fluids in the secondary bags and from the shipboard incubation bottles was near 6 (Table 1).

**Table 1.** Details of each incubation chamber, including time of deployment, mass of final and primary bag, and pH of final bag.

| Incubator Unit | Setpoint Temp (°C) | Duration (hours) | Mass of final bag (g) | Mass in primary bag (g) | pH of final bag |
| --- | --- | --- | --- | --- | --- |
| 1 (<sup>12</sup>C) | 55 | 16.55 | no data | 65 | 6.00 |
| 2 (<sup>13</sup>C) | 55 | 16.57 | 815 | 27 | 6.00 |
| 3 (<sup>12</sup>C) | 55 | 11.78 | 785 | 0 | 5.98 |
| 4 (<sup>13</sup>C) | 55 | 11.76 | 767 | 125 | 5.90 |

Both the 12 h and 16 h shipboard and incubator experiments showed ^13^C enrichment (Figure 2, Figure S2). Only the 12 h samples were sequenced to avoid heterotrophic cross-feeding from prolonged incubations. The maximum amount of 16S rRNA occurred at higher RNA densities in the ^13^C experiments versus the ^12^C controls (Figure 2) indicating that dissolved inorganic carbon (bicarbonate) was incorporated into RNA during the incubations. For the shipboard experiment, the maximum amount of 16S rRNA occurred at densities of 1.788 and 1.804 for the ^12^C control and ^13^C experiment, respectively. For the incubator experiment, maximum 16S rRNA occurred at lower RNA densities overall, with peak amounts occurring at densities of 1.778 and 1.785 for the ^12^C-control and ^13^C-experiment, respectively (Figure 2).

**Figure 2:**
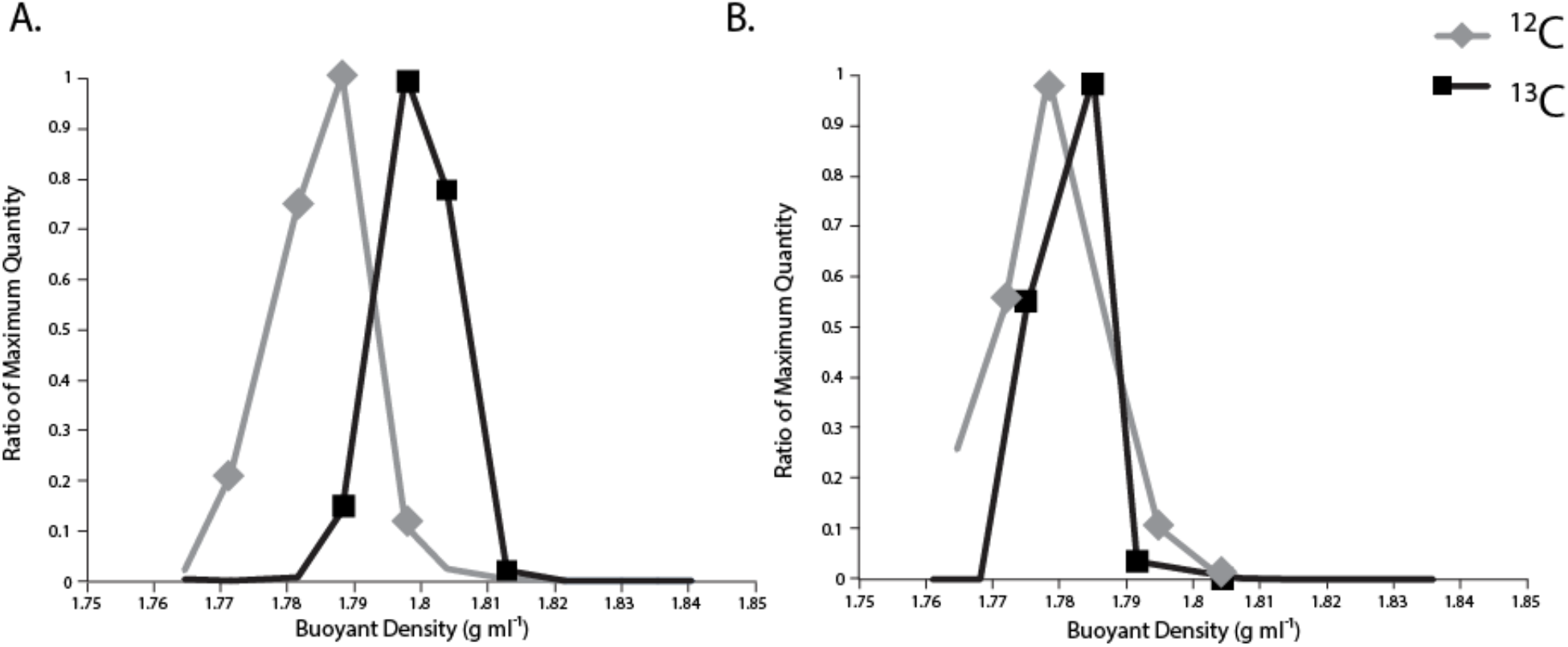
Fractionation plots of (A) Shipboard and (B) Incubator RNA-SIP experiments at 12 hours. Buoyant density (g ml-1) of each fraction is depicted on the x-axis and amount of 16S rRNA as determined by RT-qPCR is on the y-axis. Triangle and square symbols denote fractions sequenced for each experiment. Amount of 16S rRNA is displayed as the ratio of maximum quantity in order to compare between experiments.

### Taxonomic composition of RNA-SIP experiments

The taxonomic composition of the RNA-SIP experiments was determined based on the relative abundance of 16S rRNA sequences and nearly all were primarily composed (96.7% to 98.2%) of thermophilic bacteria belonging to the *Epsilonbacteraeota* (Figure 3A). The thermophilic genus *Caminibacter* was most abundant within all SIP experiments, with a relative abundance of 80.1%, 95.7% and 83.7% for the ^12^C shipboard, ^12^C incubator, and ^13^C incubator metatranscriptomes, respectively (Figure 3A). For the ^13^C shipboard experiment, *Caminibacter* was also the most abundant group but to a lesser extent, comprising 59.8% of the community, as this experiment also had a higher relative abundance of both *Nautilia* and *Hydrogenimonas* 16S rRNA sequences (Figure 3A). In the ^13^C shipboard experiment, *Nautilia* comprised 21.4% and *Hydrogenimonas* comprised 19.0% of the 16S rRNA sequences on average, indicating a different community composition in the ^13^C shipboard compared to the other experiments. *Hydrogenimonas* was more abundant in the shipboard community compared to the incubator community, where it only comprised 0.4% of the ^12^C and 8.0% of the ^13^C incubator communities on average (Figure 3A). This pattern was also observed in the taxonomic composition of the annotated transcripts (Figure 3B). While *Caminibacter* comprised close to 50% of annotated transcripts in the ^12^C shipboard, ^12^C incubator, and ^13^C incubator metatranscriptomes, transcripts classified as *Nautilia* comprised a high percentage of total annotated transcripts in all experiments (Figure 2B).

**Figure 3:**
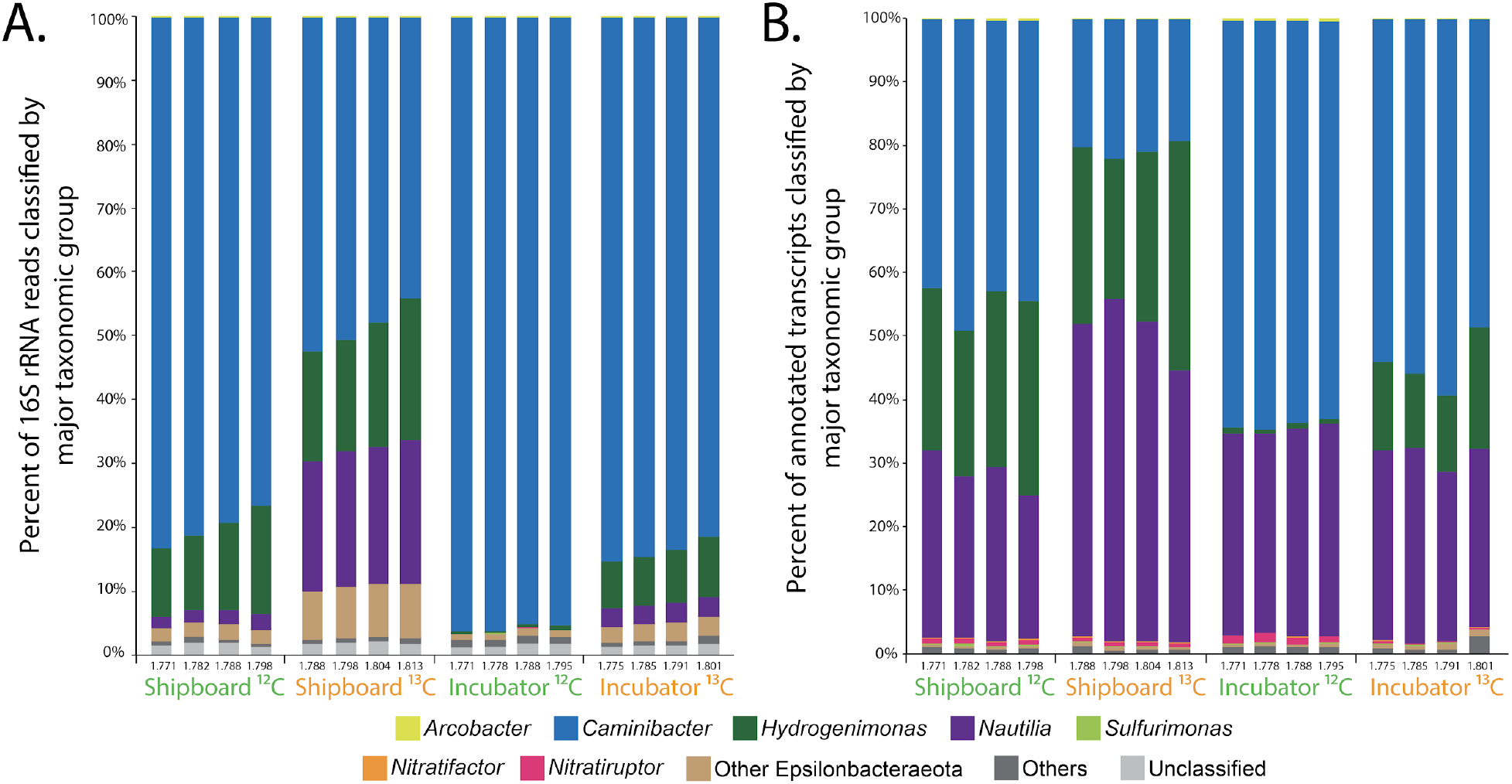
Taxonomic classification of (A) 16S rRNA reads and (B) functionally (KO) annotated non-rRNA transcripts from RNA-SIP metatranscriptomes.

Metagenome assembled genomes (MAGs) were used to examine the composition of the RNA-SIP experiments. Previously, 10 MAGs classified as thermophilic *Epsilonbacteraeota* (either *Nitratifactor* sp. or more broadly to the family *Nautiliaceae*) were identified from the Marker 33 vent metagenomic assemblies as described in Fortunato et al. (2018). A heatmap depicting mean coverage showed that these MAGs were found at various levels of coverage in the 2015 Marker 33 metagenome and were actively transcribed in the Marker 33 metatranscriptome, as well as in the RNA-SIP metatranscriptomes (Figure S3). Because of the higher coverage of the MAGs within the metagenome, patterns among the SIP experiments were masked, and therefore a second heatmap was constructed showing only mean coverage across the four SIP experiments (Figure 4). Results showed that three MAGs (Axial Epsilon Bins 1, 8, and 9) had the highest coverage across all SIP experiments. These three MAGs were broadly classified as belonging to the family *Nautiliaceae* (Figure S7). The ^13^C shipboard experiment showed additional coverage of two other MAGs (Axial Epsilon Bin 2 and 7), both also classified to the family *Nautiliaceae* (Figure 4).

**Figure 4:**
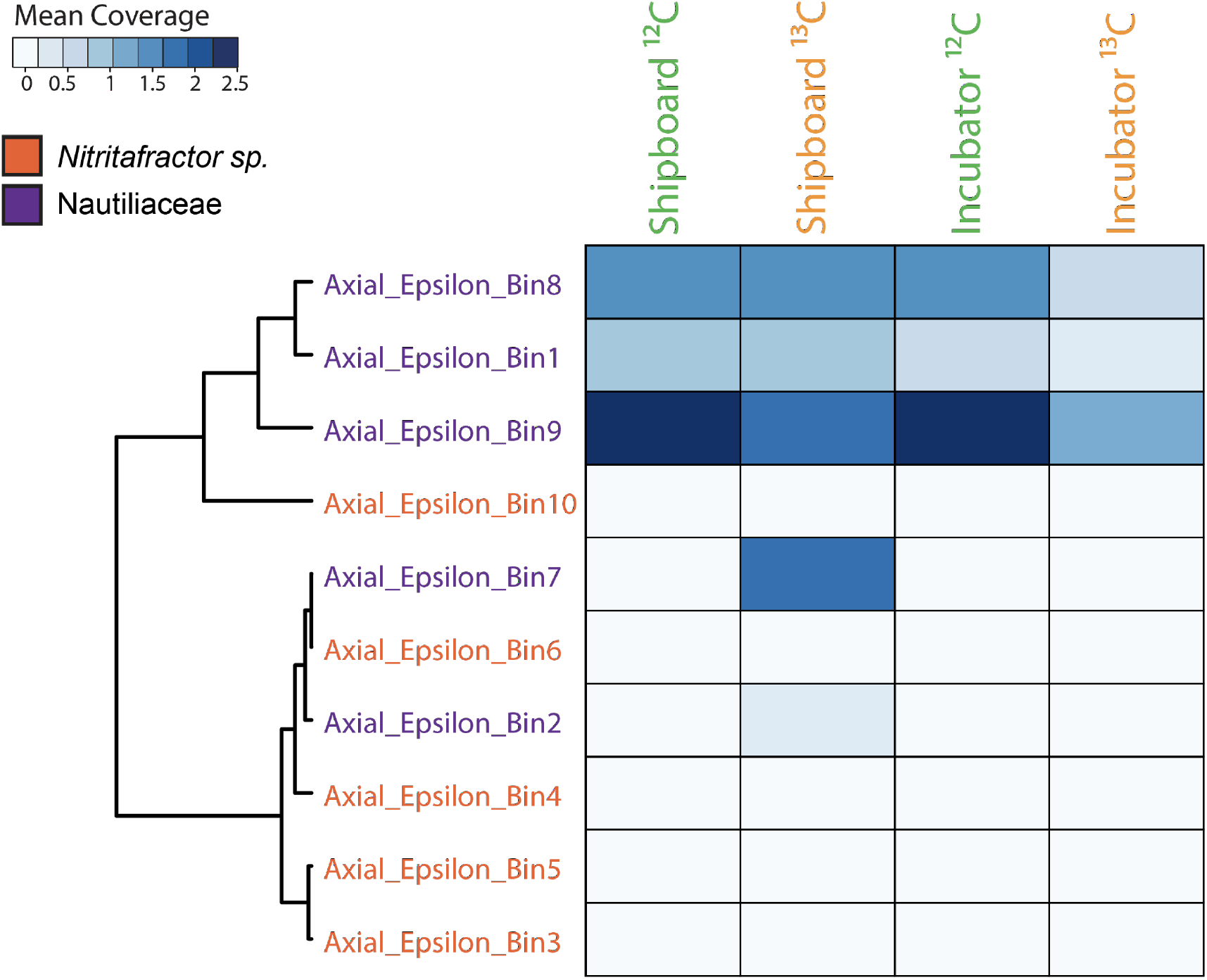
Heatmap of mean coverage across the RNA-SIP experiments of metagenome assembled genomes (MAGs) taxonomically identified as thermophilic Epsilonbacteraeota, specifically either the genus *Nitritifactor* (orange) or the family Nautiliaceae (purple) as described in Fortunato et al. 2018. Fractions from each of the four RNA-SIP experiments have been collapsed and mean coverage summed. Scale depicts range of mean coverage across MAGs.

### Determination of metabolisms within RNA-SIP experiments

Hierarchical clustering of all 16 RNA-SIP metatranscriptomes based on normalized KEGG ontology (KO) abundance of annotated transcripts showed that the ^12^C controls for the incubator and shipboard experiment clustered together, indicating functional similarity between the shipboard and incubator SIP experiments (Figure 2, Figure S4). The ^13^C experiments for the incubator and shipboard clustered separately from the ^12^C controls and from each other, with the four ^13^C shipboard metatranscriptomes forming a separate cluster (Figure S4).

When examining only the most abundant annotated transcripts expressed across all metatranscriptomes, the same clustering pattern is observed (Figure S5). The most abundant transcripts were annotated to genes related to cell growth, translational processes, and energy metabolism. The gene to which the most annotated transcripts mapped was peroxiredoxin, a gene involved in reducing oxidative stress and thus cell damage. Other highly abundant transcripts were annotated to genes for elongation factors and molecular chaperones, indications that translational machinery was active across all SIP experiments. In addition, transcripts for a key gene in the reductive TCA (rTCA) cycle, 2-oxoglutarate ferredoxin oxidoreductase, were also abundant, indicating carbon fixation was occuring (Figure S5). Additional transcripts for carbon fixation within the SIP experiments were also observed (Figure S6). As observed in the taxonomic profiles, examination of the most abundant annotated transcripts shows that the ^13^C shipboard metatranscriptome was slightly different compared to the other three experiments and clustered separately from the other metatranscriptomes (Figure S5).

Differential expression (DE) analysis was run to determine significant differences in annotated transcript abundance across the 16 RNA-SIP metatranscriptomes (Figure 5). Results showed 233 genes were significantly differentially expressed (adj. p-value < 0.01) in shipboard vs. incubator RNA-SIP libraries, with all but one being more highly expressed in shipboard experiments compared to incubator experiments (Figure 5B). Annotated transcripts with the greatest difference in expression (> 10 log2 fold change) in shipboard vs. incubator experiments included transcripts of genes related to translation, DNA replication, purine synthesis, and motility (Table S4).

**Figure 5:**
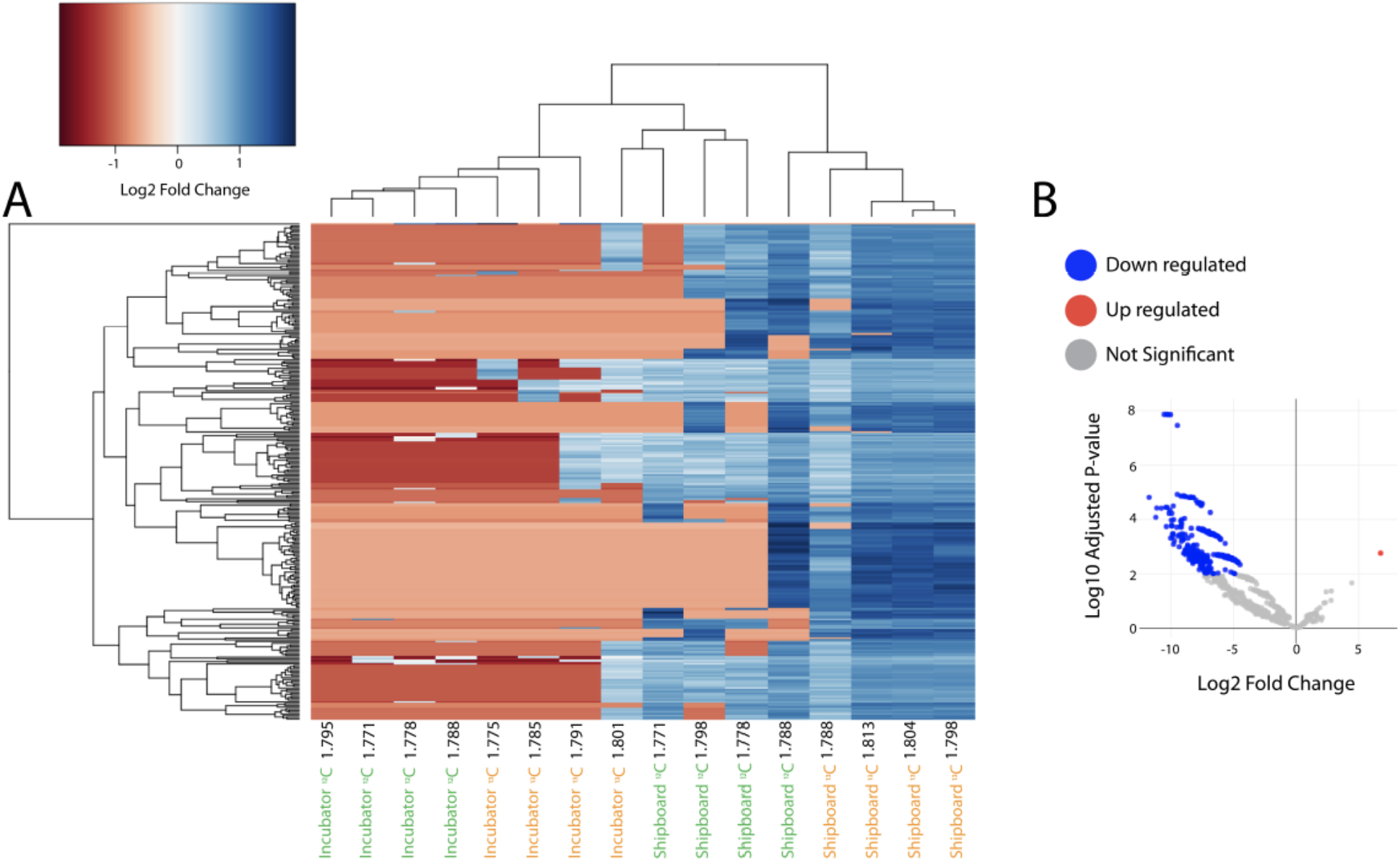
Heatmap (A) showing the 233 KO annotated genes that were differentially expressed across fractions (adj. p-value < 0.01). Volcano plot (B) of fold change in expression vs. adjusted p-value. Genes that were significantly up regulated (adj. p-value < 0.01) in the Incubator vs.Shipboard fractions are colored in red, down regulated genes are colored in blue.

The abundance of annotated transcripts involved in important metabolic processes differed between the shipboard and incubator SIP experiments (Figure 6), although DE analysis revealed that many of these differences were not significant. The metatranscriptome of diffuse fluids reflects the diversity of metabolic processes that occur at a single vent, with the presence of annotated transcripts for aerobic respiration, denitrification, methane oxidation, methanogenesis, hydrogen oxidation, sulfur reduction, and sulfur oxidation. The reduced presence of annotated transcripts observed within the RNA-SIP metatranscriptomes highlights the metabolisms active under experimental conditions. Transcripts for the mainly anaerobic process of denitrification, specifically *nirS*, *norBC*, and *nosZ*, were only observed in the shipboard experiments. Conversely, transcripts for cytochrome c oxidases, important for aerobic respiration, were only observed in one of the incubator experiments (Figure 6). Transcripts for methane metabolism differed, albeit not significantly, across experiments. Transcripts for methyl-coenzyme M reductase (*mcrA* gene), important for anaerobic methanogenesis, were only observed in the shipboard SIP experiments. Methane oxidation transcripts, however, showed an average of 1.35 log2 fold increase in incubator experiments when compared to the shipboard experiments (Figure 6). For hydrogen oxidation, transcripts annotated to genes for Group 1 Ni-Fe hydrogenases (*hydA3* and *hyaC*) were more abundant in the shipboard experiments compared to the incubator experiments (adj. p-value < 0.01). or sulfur metabolism transcripts for polysulfide and thiosulfate reduction showed a significantly higher expression (adjusted p-value < 0.01) in the shipboard compared to the incubator experiments (Figure 6).

**Figure 6:**
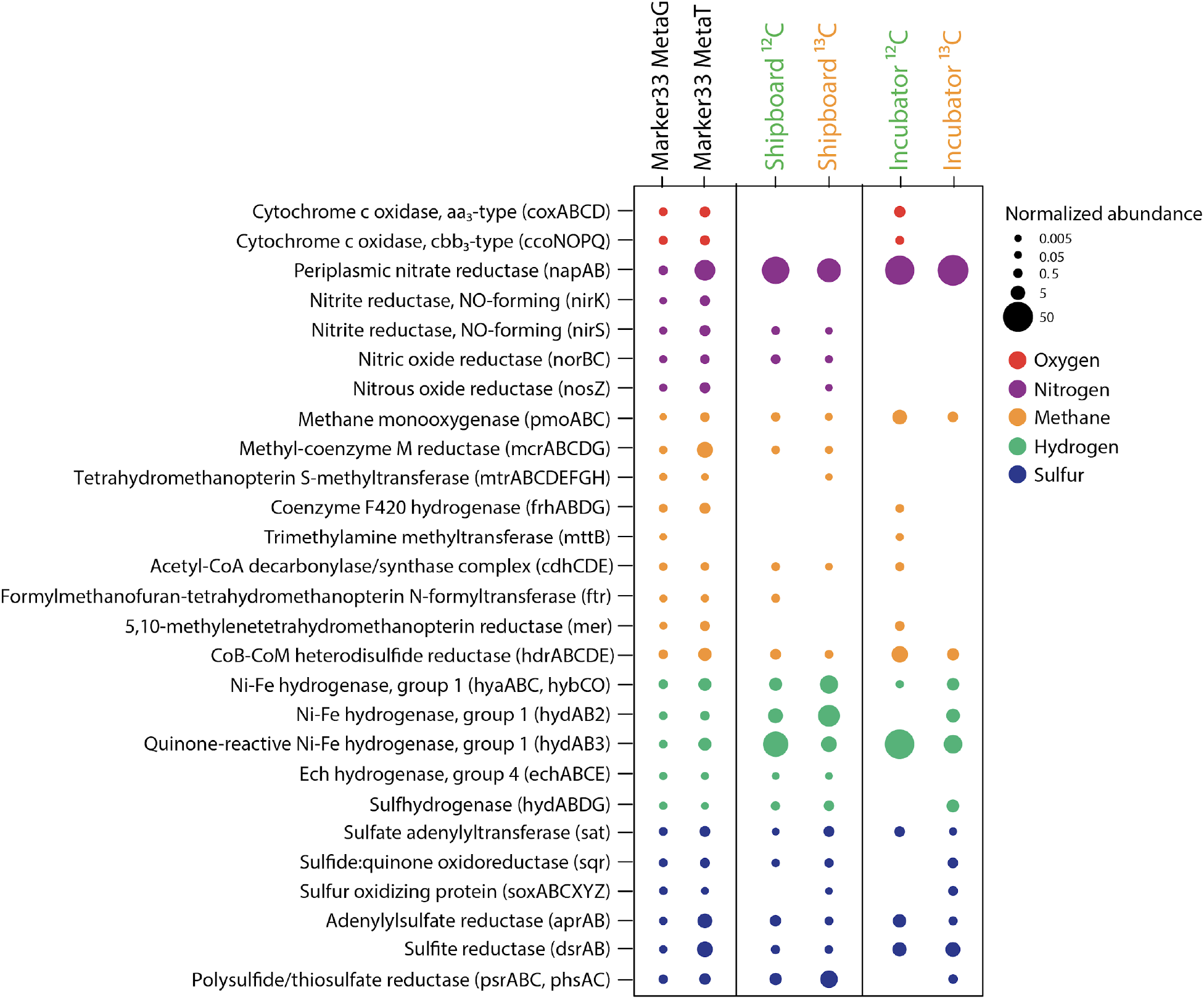
Normalized abundance of key genes and transcripts for oxygen, nitrogen, methane, hydrogen and sulfur metabolisms within the 2015 Marker 33 metagenome, metatranscriptome, and shipboard and incubator RNA-SIP experiments. Fractions from each of the four RNA-SIP experiments have been collapsed to reflect the normalized abundance of each gene in the entire experiment. Normalized abundances of metatranscriptomes were transformed to the same scale as the Marker 33 metagenome.

Because DE analysis indicated that expression of genes related to stress was significantly higher in the shipboard experiment compared to the incubator (Table S4), we further examined the expression of genes related to stress (chaperones, proteases, and other heat shock proteins) within the samples (Figure 7). An increase in abundance of transcripts annotated to heat shock chaperone genes *dnaK/dnaJ* and *GroES/GroEL* was observed in both the shipboard and incubator experiments when compared to the metatranscriptome of diffuse fluids (Figure 7). However, *dnaJ* had a significantly higher expression within the shipboard experiment (adjusted p-value < 0.01). Additionally, transcripts for the heat shock protein gene *htpX* were only expressed in the shipboard SIP metatranscriptomes. Proteases, which play an important role in protein degradation during times of stress, were in general more highly expressed in the shipboard SIP experiments when compared to the incubator. Specifically, transcripts for protease genes *clpX*, *clpP*, *ftsH*, and *hslU* all showed significantly higher expression in the shipboard experiments (adj. p-value < 0.01, Figure 7, Table S4).

**Figure 7:**
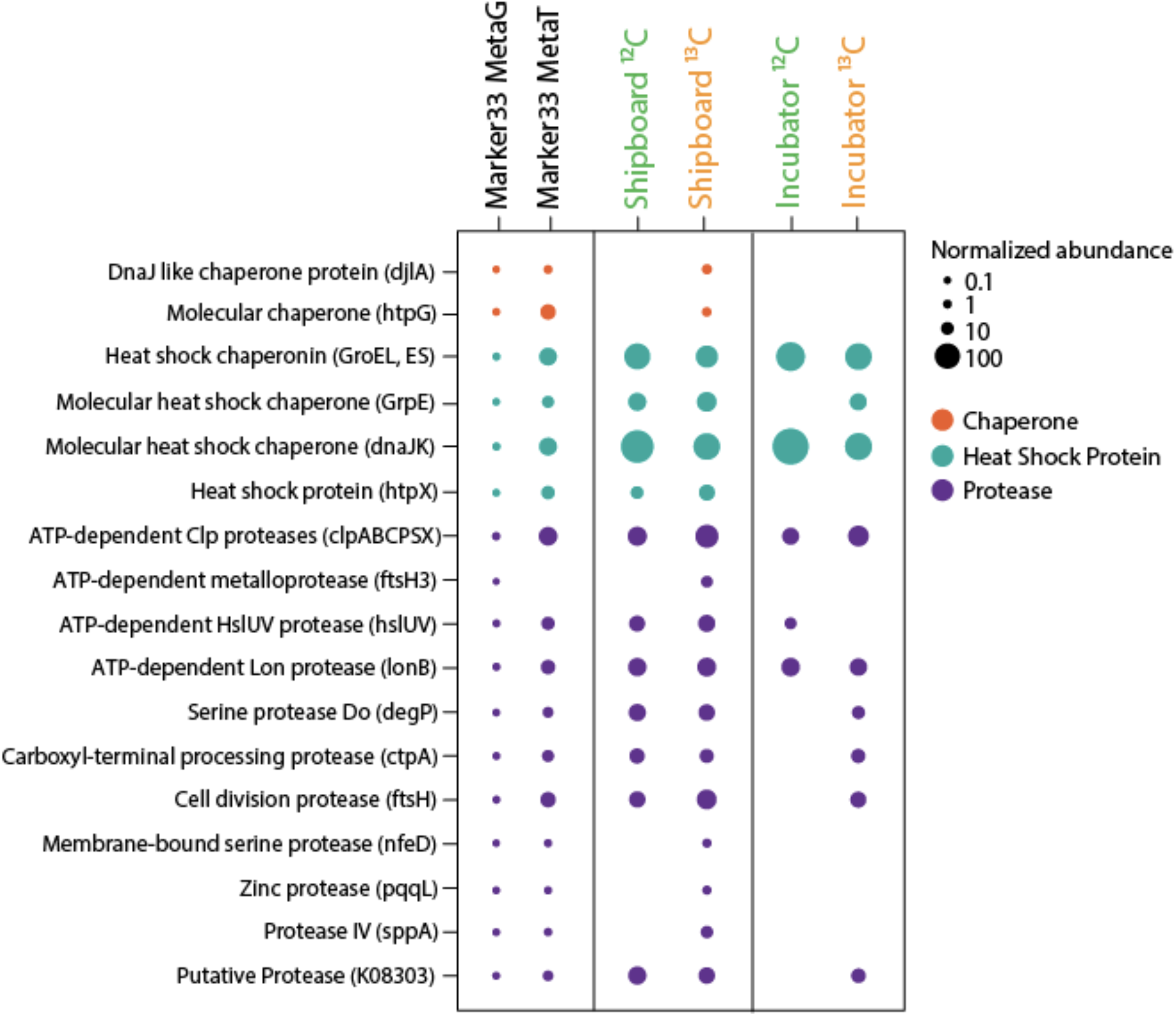
Normalized abundance genes and transcripts annotated to cell stress, including genes for protein chaperones, heat-shock proteins, and proteases within the 2015 Marker 33 metagenome, metatranscriptome, and shipboard and incubator RNA-SIP experiments. Fractions from each of the four RNA-SIP experiments have been collapsed to reflect the normalized abundance of each gene in the entire experiment. Normalized abundances of metatranscriptomes were transformed to the same scale as the Marker 33 metagenome.

## Discussion

There are extreme technical challenges to understanding microbial life in the deep sea, and results and interpretations depend heavily on the experimental approach taken. Motivated by the hypothesis that chemical reactions and microbial activity that occur in sample containers between the time of sampling and the start of an experiment will affect experimental results significantly, we designed and built an *in situ* incubator to eliminate depressurization and lag time between sampling and experiment in order to better capture the *in situ* microbial activity at deep-sea hydrothermal vents where diffusely venting fluids exit the seafloor. We successfully demonstrated the ability to study thermophilic microbes close to their seafloor and subseafloor habitats and highlighted differences between shipboard and seafloor incubations. We chose RNA Stable Isotope Probing (RNA-SIP) for the demonstration of *in situ* activity, but many different types of incubations at temperatures from ambient to at least 80°C are possible with this instrument, making it a valuable new tool for marine microbial ecology.

Marker 33 vent was chosen as the site for the seafloor incubator testing due to the consistent presence of thermophilic bacteria and archaea detected in previous studies (Fortunato et al. 2018, Huber et al. 2003, Opatkiewicz et al. 2009, Topçuoğlu et al. 2016, Stewart et al. 2019). To probe these communities, an RNA-SIP methodology coupled to mRNA sequencing was applied to examine which organisms and metabolism are responsible for autotrophy under experimental conditions that reflect those in the subseafloor (Fortunato and Huber 2016). RNA-SIP incubations must mimic the physical and chemical conditions of the environment but simulating the natural conditions of venting fluids and the subseafloor in any type of experiment is inherently challenging. For example, it may be hours before fluid collected on the seafloor can be dispensed into shipboard bottles and incubated, thus increasing the likelihood of changes to the microbial community. This time lag combined with pressure and temperature changes during transport of diffuse fluids to the surface in unpressurized vessels may also result in outgassing of key redox species such as methane, hydrogen sulfide, hydrogen, and carbon dioxide (McNichol et al 2016), death of pressure- and temperature-sensitive organisms (Fang et al 2010), or loss of oxygen to chemical reactions in the sample container. Performing experiments *in situ* on the seafloor may help ameliorate many of the biases introduced with shipboard experiments, but few direct comparisons between *in situ* versus shipboard experiments exist.

Except for location (seafloor or shipboard) and timing after fluid sampling, all other conditions were identical between experiments. The final pH of the incubations in both sets of experiments was similar to the vent fluid from Marker 33. Both the shipboard and seafloor incubator experiments showed ^13^C enrichment relative to their ^12^C control with maximum 16S rRNA occurring at higher RNA densities. However, the RNA densities of the two experiments were slightly different with peak 16S rRNA occurring at lower RNA densities overall in the incubator experiment. The reason for the lower level of enrichment in the seafloor incubator is unclear but may be due to differences in the dominant microbial genera present in each experiment or stochastic effects. The majority of the rRNA from all SIP experiments, both shipboard and incubator, was comprised of thermophilic *Epsilonbacteraeota* oxidizing hydrogen and reducing nitrate while fixing carbon, consistent with the native community present at the Marker 33 site in 2015, as well as numerous -omic surveys at diffuse vents, indicating these organisms and metabolism often dominate in the reducing, warm subseafloor habitat (Cerqueira et al 2018, Fortunato and Huber 2016, Fortunato et al 2018, McNichol et al 2018, Meier et al 2017, Olins et al 2017, Trembath-Reichert et al 2019). (Fortunato et al 2018). There was a higher percentage of rRNA classified to the genera *Hydrogenimonas* and *Nautilia* in the shipboard experiments relative to the incubator (Figure 3A) including two *Nautilia* populations only observed in the ^13^C shipboard experiment (Figure 4). Based on publicly available genomes, *Nautilia* species have a higher GC-content in their genomic DNA (average 34.8%) compared to *Caminibacter* (average 28.9%), which may account for the higher peak RNA density in the ^13^C shipboard compared to the ^13^C incubator experiment.

Differences in metabolism were apparent between the shipboard and incubator experiments and may be linked to the chemistry of the fluid at the beginning of the experiment. For example, transcripts annotated for denitrification (*nirS, norB, nosZ*) and methanogenesis (*mcrA*) were only observed in the shipboard experiments. Additionally, significantly higher expression of hydrogen oxidation transcripts (*hyaA3, hyaC,* adj. p-value < 0.01) was observed shipboard compared to the seafloor incubations. Although not significant, there was a higher abundance of transcripts annotated for methane oxidation (*pmoA*) and oxygen utilization (*cox* and *cco*) in the incubator experiments compared to shipboard (Figure 6). We hypothesize that during the lag time between sample collection and beginning the experiment shipboard, oxygen was consumed in the vent fluids by aerobic microorganisms and abiotic reactions with the high concentration of dissolved sulfide and reduced metals in the samples. Therefore, by the time the fluid was used in the shipboard incubations, there was little to no oxygen left. For the seafloor experiment, incubations of samples that were approximately 85% deep seawater (Table S1) contained oxygen at the start of the experiment and oxygen-consuming microbes grew. Aerobic oxidation of methane and sulfur species are important microbial metabolisms in hydrothermal vent plumes, as well as in many venting fluids where deep, oxygen-rich seawater mixes with the reducing vent fluids (Anantharaman et al 2016, Lesniewski et al 2012, Li et al 2014). For example, our metatranscriptomic study from multiple vent sites at Axial Seamount, including Marker 33 in 2015, showed transcription of cytochrome c oxidases and methane monooxygenase at this site, indicating these processes were occurring *in situ* in the venting fluids (Fortunato et al 2018). Additional *in situ* experiments focused on assessing the metatranscriptome of the incubated vent fluid over a shorter time scale might resolve an initial aerobic stage from a later anaerobic stage and capture some of the dynamic spatial variability in microbial activity around diffuse vent sites.

In addition to differences in microbial metabolism, we found significantly higher expression of transcripts annotated to heat-shock proteins, proteases, and chaperones in the shipboard experiments compared to the incubator, which may indicate that the shipboard microbial community was under more thermal stress (Stewart et al 2012). Chaperones can aid in protein folding and prevent protein denaturation that occurs during environmental stress (Stewart et al 2012, Susin et al 2006). Transcripts for chaperone encoding genes were expressed in both shipboard and incubator experiments (Figure 7), an indication that experimental incubations, be it on the seafloor or shipboard, enact some stress on microbial communities. However, transcripts annotated as proteases and heat shock proteins were significantly more abundant in the shipboard experiments (adj. p-value < 0.01), particularly in the ^13^C experiment (Figure 7).

The increased environmental stress could be due to transport to atmospheric pressure, manipulation of fluid into glass bottles, or any number of differences that occur when carrying out incubations shipboard as compared to incubating the fluid *in situ* on the seafloor. Another possibility is that the incubations were performed at temperatures near the optimal growth temperatures of *Caminibacter* (55-60°C, Alain et al. 2002, Miroshnichenko et al. 2004, Voordeckers et al. 2005), *Nautilia* (53-60°C, Miroshnichenko et al. 2002, Alain et al. 2009, Perez-Rodriguez et al. 2010), and *Hydrogenimonas* (55°C, Takai et al. 2004), which may induce transcription of thermal stress proteins in these organisms. Growth at pressures found at deep-sea vents increased the optimal growth temperature (Jannasch et al. 1992, Pledger et al. 1994, Holden and Baross 1995) and raised the thermal induction temperature (Holden and Baross 1995) in hyperthermophilic archaea. Therefore, *in situ* incubation of vent fluids in this study may similarly ameliorate thermal stress in *Epsilonbacteraeota* relative to shipboard incubations.

In conclusion, this study showed the effects of depressurization and sample processing delay using a new *in situ* incubator instrument to carry out RNA-SIP experiments in situ on the seafloor. The taxonomic and functional gene differences observed between shipboard and incubator experiments were likely due to slight differences in the chemistry of the fluid at the start of the experiment, and more specifically, the availability of oxygen in the incubator experiment. Microbial populations were also more stressed in shipboard experiments. Although the shipboard and incubator experiments were similar, the slight differences between the two suggest that use of a seafloor incubator may give a more accurate account of the microbial metabolic processes occurring within diffusely venting fluids due to reduced lag time, depressurization, and stress, as well as limiting both abiotic and biotic reactions that modify the chemistry of the fluids during transport to the ship.

Use of instrumentation like the seafloor incubator is an important step in understanding and constraining the roles microbial communities play in the deep ocean, with potential applications well beyond those described here. The incubator can collect seawater, cold seep fluids, or vent fluids and their associated microbial communities and immediately amend the fluids while keeping them at *in situ* pressure and a controlled temperature before filtering and preserving the microbial biomass. Future experiments with the incubator will focus on performing quantitative time series measurements of microbial, viral, and geochemical activity for various biogeochemical processes, as well as nutrient amendment experiments to measure the effect of substrate concentration on reaction rates, chemical signatures, and microbial and viral community composition and function. Thus, our study expands our understanding of the activities of natural microbial assemblages under near-native conditions at deep-sea hydrothermal vents and allows for future deployments to better constrain marine microbial biogeochemistry in the ocean.

## Acknowledgements

This work was funded by the Gordon and Betty Moore Foundation Grant GBMF3297, the NSF Center for Dark Energy Biosphere Investigations (C-DEBI) (OCE-0939564), NOAA/PMEL, contribution number 5182, and Joint Institute for the Study of the Atmosphere and Ocean (JISAO) under NOAA Cooperative Agreement NA15OAR4320063, contribution number 2020-1113. We thank the captains and crews of the R/V *Thomas G. Thompson* and R/V Brown, and in particular the ROV *Jason* II group for assistance with incubator integration on the vehicle. Andra Bobbit, Bill Chadwick, Kevin Roe, Susan Merle, and Ryan Wells provided critical support at sea. We also thank Paula Pelayo who helped with RT-qPCR assay development. NOAA/PMEL supported this work with ship time in 2014 and through funding to the Earth Ocean Interactions group. NSF provided shiptime for the 2015 expedition through OCE 1546695 to DAB and OCE-1547004 to JFH.

## References

Akerman NH, Butterfield DA, Huber JA (2013). Phylogenetic diversity and functional gene patterns of sulfur-oxidizing subseafloor Epsilonproteobacteria in diffuse hydrothermal vent fluids. Front Microbiol 4.

Alain K, Querellou J, Lesongeur F, Pignet P, Crassous P, Raguenes G, Cueff V, Cambon-Bonavita MA (2002). *Caminibacter hydrogeniphilus* gen. nov., sp. nov., a novel thermophilic, hydrogen-oxidizing bacterium isolated from an East Pacific Rise hydrothermal vent. Int J Syst Evol Microbiol 52: 1317–1323.

Alain K, Callac N, Guegan M, Lesongeur F, Crassous P, Cambon-Bonavita MA, Querellou J, Prieur D (2009). *Nautilia abyssi* sp. nov., a thermophilic, chemolithoautotrophic, sulfur-reducing bacterium isolated from an East Pacific Rise hydrothermal vent. Int J Syst Evol Microbiol 59: 1310–1315.

Anantharaman K, Breier JA, Dick GJ (2016). Metagenomic resolution of microbial functions in deep-sea hydrothermal plumes across the Eastern Lau Spreading Center. Isme J 10: 225–239.

Bourbonnais A, Juniper S, Butterfield D, Devol A, Kuypers M, Lavik G et al (2012). Activity and abundance of denitrifying bacteria in the subsurface biosphere of diffuse hydrothermal vents of the Juan de Fuca Ridge. Biogeosciences discussions 9: 4661–4678.

Butterfield DA, Roe KK, Lilley MD, Huber JA, Baross JA, Embley RW et al (2004). Mixing, Reaction and Microbial Activity in the Sub-Seafloor Revealed by Temporal and Spatial Variation in Diffuse Flow Vents at Axial Volcano. The Subseafloor Biosphere at Mid-Ocean Ridges. American Geophysical Union. pp 269–289.

Campbell BJ, Smith JL, Hanson TE, Klotz MG, Stein LY, Lee CK et al (2009). Adaptations to Submarine Hydrothermal Environments Exemplified by the Genome of Nautilia profundicola. Plos Genetics 5.

Cario A, Oliver GC, Rogers KL (2019). Exploring the Deep Marine Biosphere: Challenges, Innovations, and Opportunities. Frontiers in Earth Science 7.

Cerqueira T, Barroso C, Froufe H, Egas C, Bettencourt R (2018). Metagenomic signatures of microbial communities in deep-sea hydrothermal sediments of Azores vent fields. Microb Ecol 76: 387–403.

Corliss J, Ballard R (1977). Oases of life in the cold abyss. National Geographic 152: 441–453.

Darling AE, Jospin G, Lowe E, Matsen FAIV, Bik HM, Eisen JA (2014). PhyloSift: phylogenetic analysis of genomes and metagenomes. PeerJ 2: e243.

Edgcomb VP, Taylor C, Pachiadaki MG, Honjo S, Engstrom I, Yakimov M (2016). Comparison of Niskin vs. in situ approaches for analysis of gene expression in deep Mediterranean Sea water samples. Deep Sea Research Part II: Topical Studies in Oceanography 129: 213–222.

Eren AM, Esen ÖC, Quince C, Vineis JH, Morrison HG, Sogin ML et al (2015). Anvi’o: an advanced analysis and visualization platform for ‘omics data. PeerJ 3: e1319.

Fang J, Zhang L, Bazylinski DA (2010). Deep-sea piezosphere and piezophiles: geomicrobiology and biogeochemistry. Trends Microbiol 18: 413–422.

Fortunato CS, Huber JA (2016). Coupled RNA-SIP and metatranscriptomics of active chemolithoautotrophic communities at a deep-sea hydrothermal vent. Isme J 10: 1925–1938.

Fortunato CS, Larson B, Butterfield DA, Huber JA (2018). Spatially distinct, temporally stable microbial populations mediate biogeochemical cycling at and below the seafloor in hydrothermal vent fluids. Environ Microbiol 20: 769–784.

Frank KL, Rogers DR, Olins HC, Vidoudez C, Girguis PR (2013). Characterizing the distribution and rates of microbial sulfate reduction at Middle Valley hydrothermal vents. The ISME Journal 7: 1391–1401.

Frank KL, Rogers KL, Rogers DR, Johnston DT, Girguis PR (2015). Key Factors Influencing Rates of Heterotrophic Sulfate Reduction in Active Seafloor Hydrothermal Massive Sulfide Deposits. Front Microbiol 6.

Galambos D, Anderson RE, Reveillaud J, Huber JA (2019). Genome-resolved metagenomics and metatranscriptomics reveal niche differentiation in functionally redundant microbial communities at deep-sea hydrothermal vents. Environ Microbiol 21: 4395–4410.

Holden JF, Baross JA (1995). Enhanced thermotolerance by hydrostatic pressure in the deep-sea hyperthermpphile *Pyrococcus* strain ES4. FEMS Microbiol Ecol 18: 27–34.

Huber JA, Butterfield DA, Baross JA (2003). Bacterial diversity in a subseafloor habitat following a deep-sea volcanic eruption. Fems Microbiol Ecol 43: 393–409.

Huber JA, Mark Welch D, Morrison HG, Huse SM, Neal PR, Butterfield DA et al (2007). Microbial population structures in the deep marine biosphere. Science 318: 97–100.

Jacobson M, Sylvan J, Edwards K (2014). Extracellular Enzyme Activity and Microbial Diversity Measured on Seafloor Exposed Basalts from Loihi Seamount Indicate the Importance of Basalts to Global Biogeochemical Cycling. Appl Environ Microb 80.

Jannasch HW, Wirsen CO, Winget CL (1973). A bacteriological pressure-retaining deep-sea sampler and culture vessel. Deep Sea Research and Oceanographic Abstracts 20: 661–664.

Jannasch HW, Mottl MJ (1985). Geomicrobiology of Deep-Sea Hydrothermal Vents. Science 229: 717–725.

Jannasch HW, Wirsen CO, Molyneaux SJ, Langworthy TA (1992). Comparative physiological studies on hyperthermophilic archaea isolated from deep-sea hydrothermal vents. Appl Environ Microbiol 58: 3472–3481.

Karl DM (1995). The microbiology of deep-sea hydrothermal vents. CRC Press.

Kucukural, A, Yukselen, O, Ozata, DM, Moore, MJ, Garber, M (2019). DEBrowser: interactive differential expression analysis and visualization tool for count data. BMC Genomics 20: 6

Langmead B, Salzberg SL (2012). Fast gapped-read alignment with Bowtie 2. Nat Meth 9: 357–359.

Lesniewski RA, Jain S, Anantharaman K, Schloss PD, Dick GJ (2012). The metatranscriptome of a deep-sea hydrothermal plume is dominated by water column methanotrophs and lithotrophs. Isme J 6: 2257–2268.

Li H, Handsaker B, Wysoker A, Fennell T, Ruan J, Homer N et al (2009). The Sequence Alignment/Map format and SAMtools. Bioinformatics (Oxford, England) 25: 2078–2079.

Li M, Jain S, Baker BJ, Taylor C, Dick GJ (2014). Novel hydrocarbon monooxygenase genes in the metatranscriptome of a natural deep-sea hydrocarbon plume. Environ Microbiol 16: 60–71.

Lippsett L (2014). Bringing a Lab to the Seafloor: New device probes deep-sea microbial life. Oceanus magazine. Woods Hole Oceanographic Institute.

McNichol J, Sylva SP, Thomas F, Taylor CD, Sievert SM, Seewald JS (2016). Assessing microbial processes in deep-sea hydrothermal systems by incubation at in situ temperature and pressure. Deep Sea Research Part I: Oceanographic Research Papers 115: 221–232.

McNichol J, Stryhanyuk H, Sylva SP, Thomas F, Musat N, Seewald JS et al (2018). Primary productivity below the seafloor at deep-sea hot springs. Proceedings of the National Academy of Sciences 115: 6756.

McQuillan JS, Robidart JC (2017). Molecular-biological sensing in aquatic environments: recent developments and emerging capabilities. Current opinion in biotechnology 45: 43–50.

Medina LE, Taylor CD, Pachiadaki MG, Henríquez-Castillo C, Ulloa O, Edgcomb VP (2017). A Review of Protist Grazing Below the Photic Zone Emphasizing Studies of Oxygen-Depleted Water Columns and Recent Applications of In situ Approaches. Frontiers in Marine Science 4.

Meier DV, Pjevac P, Bach W, Hourdez S, Girguis PR, Vidoudez C et al (2017). Niche partitioning of diverse sulfur-oxidizing bacteria at hydrothermal vents. Isme J 11: 1545–1558.

Miroshnichenko ML, Kostrikina NA, L’Haridon S, Jeanthon C, Hippe H, Stackebrandt E, Bonch-Osmolovskaya EA (2002). *Nautilia lithotrophica* gen. nov., sp.nov., a thermophilic sulfur-reducing ∊-proteobacterium isolated from a deep-sea hydrothermal vent. Int J Syst Evol Microbiol 52: 1299–1304.

Miroshnichenko ML, L’Haridon S, Schumann P, Spring S, Bonch-Osmolovskaya EA, Jeanthon C, Stackebrandt E (2004). *Caminibacter profundus* sp. nov., a novel thermophile of *Nautiliales* ord. nov. within the class ‘*Epsilonproteobacteria*’, isolated from a deep-sea hydrothermal vent. Int J Syst Evol Microbiol 54: 41–45.

Olins HC, Rogers DR, Frank KL, Vidoudez C, Girguis PR (2013). Assessing the influence of physical, geochemical and biological factors on anaerobic microbial primary productivity within hydrothermal vent chimneys. Geobiology 11: 279–293.

Olins HC, Rogers DR, Preston C, Ussler W, Pargett D, Jensen S et al (2017). Co-registered Geochemistry and Metatranscriptomics Reveal Unexpected Distributions of Microbial Activity within a Hydrothermal Vent Field. Front Microbiol 8.

Opatkiewicz AD, Butterfield DA, Baross JA (2009). Individual hydrothermal vents at Axial Seamount harbor distinct subseafloor microbial communities. Fems Microbiol Ecol 70: 413–424.

Orcutt BN, Sylvan JB, Rogers D, Delaney J, Lee RW, Girguis PR (2015). Carbon fixation by basalt-hosted microbial communities. Front Microbiol 6: 904.

Ottesen EA (2016). Probing the living ocean with ecogenomic sensors. Current Opinion in Microbiology 31: 132–139.

Pachiadaki M, Rédou Creff V, Beaudoin D, Burgaud G, Edgcomb V (2016). Fungal and Prokaryotic Activities in the Marine Subsurface Biosphere at Peru Margin and Canterbury Basin Inferred from RNA-Based Analyses and Microscopy. Front Microbiol 7.

Parks DH, Imelfort M, Skennerton CT, Hugenholtz P, Tyson GW (2015). CheckM: assessing the quality of microbial genomes recovered from isolates, single cells, and metagenomes. Genome Research 25: 1043–1055.

Perez-Rodriguez I, Ricci J, Voordeckers JW, Starovoytov V, Vetriani C (2010). *Nautilia nitratireducens* sp. nov., a thermophilic, anaerobic, chemosynthetic, nitrate-ammonifying bacterium isolated from a deep-sea hydrothermal vent. Int J Syst Evol Microbiol 60: 1182–1186.

Perner M, Bach W, Hentscher M, Koschinsky A, Garbe-Schonberg D, Streit WR et al (2009). Short-term microbial and physico-chemical variability in low-temperature hydrothermal fluids near 5 degrees S on the Mid-Atlantic Ridge. Environ Microbiol 11: 2526–2541.

Perner M, Petersen JM, Zielinski F, Gennerich H-H, Seifert R (2010). Geochemical constraints on the diversity and activity of H-2-oxidizing microorganisms in diffuse hydrothermal fluids from a basalt- and an ultramafic-hosted vent. Fems Microbiol Ecol 74: 55–71.

Pledger RJ, Crump BC, Baross JA (1994) A barophilic response by two hyperthermophilic, hydrothermal vent Archaea: an upward shift in the optimal temperature and acceleration of growth rate at supra-optimal temperatures by elevated pressure. FEMS Microbiol Ecol 14: 233–242.

R-Development-Core-Team (2011). R: A language and environment for statistical computing. R Foundation for Statistical Computing.

Reveillaud J, Reddington E, McDermott J, Algar C, Meyer JL, Sylva S et al (2016). Subseafloor microbial communities in hydrogen-rich vent fluids from hydrothermal systems along the Mid-Cayman Rise. Environ Microbiol 18: 1970–1987.

Ritchie ME, Phipson B, Wu D, Hu Y, Law CW, Shi W, Smyth GK (2015). limma powers differential expression analyses for RNA-sequencing and microarray studies. Nucleic Acids Res 43(7): e47

Scholin CA, J. Birch, S. Jensen, R. Marin III, E. Massion, D. Pargett, C. Preston, B. Roman, and W. Ussler III (2018). The Quest to Develop Ecogenomic Sensors: A 25-Year History of the Environmental Sample Processor (ESP) as a Case Study. Oceanography 30: 100–113.

Sievert SM, Vetriani C (2012). Chemoautotrophy at Deep-Sea Vents Past, Present, and Future. Oceanography 25: 218–233.

Stewart FJ, Dalsgaard T, Young CR, Thamdrup B, Revsbech NP, et al. (2012) Experimental Incubations Elicit Profound Changes in Community Transcription in OMZ Bacterioplankton. PLOS ONE 7(5): e37118. https://doi.org/10.1371/journal.pone.0037118

Stewart LC, Algar CK, Fortunato CS, Larson BI, Vallino JJ, Huber JA et al (2019). Fluid geochemistry, local hydrology, and metabolic activity define methanogen community size and composition in deep-sea hydrothermal vents. The ISME Journal.

Susin MF, Baldini RL, Gueiros-Filho F, Gomes SL (2006). GroES/GroEL and DnaK/DnaJ have distinct roles in stress responses and during cell cycle progression in Caulobacter crescentus. J Bacteriol 188: 8044–8053.

Sweetman AK, Smith CR, Shulse CN, Maillot B, Lindh M, Church MJ et al (2019). Key role of bacteria in the short-term cycling of carbon at the abyssal seafloor in a low particulate organic carbon flux region of the eastern Pacific Ocean. Limnol Oceanogr 64: 694–713.

Takahashi S, Tomita J, Nishioka K, Hisada T, Nishijima M (2014). Development of a prokaryotic universal primer for simultaneous analysis of Bacteria and Archaea using next-generation sequencing. Plos One 9: e105592.

Takai K, Nealson KH, Horikoshi K (2004). *Hydrogenimonas thermophila* gen. nov., sp. nov., a novel thermophilic, hydrogen-oxidizing chemolithoautotroph within the ∊-*Proteobacteria*, isolated from a black smoker in a Central Indian Ridge hydrothermal field. Int J Syst Evol Microbiol 54: 25–32.

Taylor C, Howes BL, Doherty KW (1993). Automated instrumentation for time-series measurement of primary production and nutrient status in production platform-accessible environments. Marine Technology Society Journal 27: 32–44.

Taylor CD, Molongoski JJ, Lohrenz SE (1983). Instrumentation for the measurement of phytoplankton production 1. Limnol Oceanogr 28: 781–787.

Taylor CD, Doherty KW (1990). Submersible Incubation Device (SID), autonomous instrumentation for the in situ measurement of primary production and other microbial rate processes. Deep Sea Research Part A Oceanographic Research Papers 37: 343–358.

Trembath-Reichert E, Butterfield DA, Huber JA (2019). Active subseafloor microbial communities from Mariana back-arc venting fluids share metabolic strategies across different thermal niches and taxa. The ISME Journal 13: 2264–2279.

Topcuoglu BD, Stewart LC, Morrison HG, Butterfield DA, Huber JA, Holden JF (2016). Hydrogen limitation and syntrophic growth among natural assemblages of thermophilic methanogens at deep-sea hydrothermal vents. Front Microbiol 7: 1240.

Tuttle J, Wirsen C, Jannasch H (1983). Microbial activities in the emitted hydrothermal waters of the Galapagos Rift vents. Marine Biology 73: 293–299.

Ussler W, Preston C, Tavormina P, Pargett D, Jensen S, Roman B et al (2013). Autonomous Application of Quantitative PCR in the Deep Sea: In Situ Surveys of Aerobic Methanotrophs Using the Deep-Sea Environmental Sample Processor. Environmental Science & Technology 47: 9339–9346.

Ver Eecke HC, Butterfield DA, Huber JA, Lilley MD, Olson EJ, Roe KK et al (2012). Hydrogen-limited growth of hyperthermophilic methanogens at deep-sea hydrothermal vents. P Natl Acad Sci USA 109: 13674–13679.

Voordeckers JW, Starovoytov V, Vetriani C (2005). *Caminibacter mediatlanticus* sp. nov., a thermophilic, chemolithoautotrophic, nitrate-ammonifying bacterium isolated from a deep-sea hydrothermal vent on the Mid-Atlantic Ridge. Int J Syst Evol Microbiol 55: 773–779.

Wirsen CO, Tuttle J, Jannasch H (1986). Activities of sulfur-oxidizing bacteria at the 21 N East Pacific Rise vent site. Marine Biology 92: 449–456.

Witte U, Wenzhöfer F, Sommer S, Boetius A, Heinz P, Aberle N et al (2003). In situ experimental evidence of the fate of a phytodetritus pulse at the abyssal sea floor. Nature 424: 763–766.

ZoBell CE (1941). Apparatus for collecting water samples from different depths for bacteriological analysis. J mar Res 4: 173–188.

