## Supplemental Methods, Tables, and Figures for "Seafloor incubation experiment with deep-sea hydrothermal vent fluid reveals effect of pressure and lag time on autotrophic microbial communities"

***Supplemental Material:***

### Supplemental Methods:

*SIP RNA Extraction, Density Fractionation, and qRT-PCR*

RNA from the incubator and shipboard SIP experiments was extracted using the mirVana miRNA isolation kit with added bead-beating step using RNA PowerSoil beads (Fortunato and Huber 2016). Extracted RNA (100 µl) was then DNase treated using the Turbo-DNase kit. Gradient preparation, isopycnic centrifugation, and fractionation were performed as described in Fortunato and Huber (2016). Each gradient was fractionated into 12 tubes of approximately 410 µl and the refractive index of each was measured to determine density. RNA was then precipitated using ice-cold isopropanol and washed with 70% ethanol as described in Lueders (2010). RNA concentration of each fraction was determined using the RiboGreen quantification kit and a Gemini XPS plate reader. RT-qPCR was completed using the KAPA SYBR FAST One-Step qRT-PCR kit and universal primers Pro341F and Pro805R (Takahashi et al 2014). Primers were optimized to a final concentration of 0.2 µM and 0.4 µM for the forward and reverse primer respectively. *E. coli* ribosomal RNA was used to construct a standard curve. RT-qPCR results were used to determine the ratio of maximum quantity of 16S rRNA within each set of fractions, that is the ratio of 16S rRNA in a given fraction to the maximum copy number across all fractions.

*SIP metatranscriptomic library preparation*

Using results from RT-qPCR, four fractions from each of the $^{12}C$ and $^{13}C$ samples from the shipboard and incubator samples were sequenced, including fractions with the peak number of 16S rRNA genes and fractions on either side of the peak, for a total of 16 metatranscriptomic libraries. For each SIP metatranscriptomic library, double stranded cDNA was constructed using SuperScript III First-strand synthesis system and mRNA second strand synthesis module. Double stranded cDNA was sheared to a fragment size of 275bp using a Covaris S-series sonicator. SIP metatranscriptomic library construction was completed using the Ovation Ultralow Library DR multiplex system. following manufacturer's instructions. Sequencing was performed on an Illumina NextSeq 500 at the W.M. Keck sequencing facility at the Marine Biological Laboratory. All libraries were paired-end, with a 30 bp overlap, resulting in an average merged read length of 275 bp.

*Metagenomic and metatranscriptomic library preparation*

The 47mm flat filters were cut in half with a sterile razor, with half used for DNA and half used for RNA extraction. RNA was extracted using the mirVana miRNA isolation kit with added bead-beating step using RNA PowerSoil beads. A total volume of 100 µl was extracted and was then DNase treated using the Turbo-DNase kit (Ambion), purified, and concentrated using the RNAeasy MinElute kit. Ribosomal RNA removal, cDNA synthesis, and metatranscriptomic library preparation was carried out using the Ovation Complete Prokaryotic RNA-Seq DR multiplex system following manufacturer instructions. Prior to library construction, cDNA was sheared to a fragment size of 275 bp using a Covaris S-series sonicator. For DNA extraction, the DNA filter was first rinsed with sterile Phosphate Buffered Saline (PBS) to remove RNAlater and then was extracted using a phenol-chloroform method adapted from Crump et al. (2003) and Zhou et al. (1996). DNA was then sheared to a fragment size of 275 bp using a Covaris S-series sonicator. Metagenomic library construction was completed using the Ovation Ultralow Library DR multiplex system (Nugen) following manufacturer instructions. Metagenomic and metatranscriptomic sequencing was performed on an Illumina Nextseq 500 at the W.M. Keck sequencing facility at the Marine Biological Laboratory. All libraries were paired-end, with a 30 bp overlap, resulting in an average merged read length of 275 bp.

*Analysis of Marker 33 metagenome, metatranscriptome, and RNA-SIP metatranscriptomes:*

To identify ribosomal RNA reads, the Marker 33 metatranscriptome and RNA-SIP metatranscriptomes were mapped to SILVA SSU and LSU databases (release 123, Pruesse et al 2007) using Bowtie2 with a local alignment and default settings (v2.0.0-beta5 Langmead and Salzberg 2012). For the Marker 33 metatranscriptome, identified rRNA reads were removed as described in Fortunato et al. (2018). For the RNA-SIP metatranscriptomes, 16S rRNA reads were taxonomically identified via MOTHUR (v1.37, Schloss et al 2009) using the Greengenes 16S rRNA taxonomic database (August 2013 release, McDonald et al 2012).

The Marker 33 metagenome was assembled as described in Fortunato et al. (2018) and contigs were submitted to the DOE Joint Genome Institute's Integrated Microbial Genome Metagenomic Expert Review (IMG/MER) annotation pipeline for Open Reading Frame (ORF) identification and functional and taxonomic annotation (Markowitz et al 2012). Identified ORFs from the Marker 33 metagenome were annotated against the KEGG orthology (KO) database as described in Fortunato et al (2018). Metagenomic reads were mapped to each ORF using Bowtie2 (v2.0.0-beta5 Langmead and Salzberg 2012), with end to end alignment. For the Marker 33 metatranscriptome as well as all 16 RNA-SIP metatranscriptomes, non-rRNA transcripts were mapped to ORFs identified in the Marker 33 metagenome. After mapping, abundances for each ORF were normalized to gene or transcript length. Metagenomic read abundance was normalized to the number of Reads per Kilobase per Genome (RPKG). Number of genomes per metagenome was estimated using hits to the single-copy gene, DNA-directed RNA polymerase beta subunit gene (*rpoB*). The Marker 33 metatranscriptome and all RNA-SIP metatranscriptomes were normalized to the number of Transcripts Per Million reads (TPM). Normalized KO abundances were used for bubble plot construction, hierarchical clustering, and heatmap generation. Hierarchical clustering of samples (average-linkage method) was carried out using the statistical program R (v3.3.2, R-Development-Core-Team 2011) and the R package *pvclust*. Heatmaps were constructed in R using the package *heatmap3*.

To determine taxonomy of non-rRNA transcripts as shown in Figure 3, the RNA-SIP metatranscriptomes were assembled using CLC Genomics Workbench (v 7.0) using default settings and submitted to IMG/MER for ORF identification. The taxonomy for each ORF was determined using the Phylogenetic Distribution tool in IMG, part of the IMG/MER annotation pipeline. To determine taxonomic abundance, the non-rRNA reads were then mapped to each ORF using Bowtie2 (v2.0.0-beta5 Langmead and Salzberg 2012).

**References:**


Crump BC, Kling GW, Bahr M, Hobbie JE (2003). Bacterioplankton community shifts in an arctic lake correlate with seasonal changes in organic matter source. *Appl Environ Microb* **69:** 2253-2268.

Fortunato CS, Huber JA (2016). Coupled RNA-SIP and metatranscriptomics of active chemolithoautotrophic communities at a deep-sea hydrothermal vent. *Isme J* **10:** 1925-1938

Fortunato CS, Larson B, Butterfield DA, Huber JA (2018). Spatially distinct, temporally stable microbial populations mediate biogeochemical cycling at and below the seafloor in hydrothermal vent fluids. *Environ Microbiol* **20:** 769-784.

Lueders T (2010). Stable Isotope Probing of Hydrocarbon-Degraders. In: Timmis K (ed). *Handbook of Hydrocarbon and Lipid Microbiology*. Springer Berlin Heidelberg. pp 4011-4026.

Langmead B, Salzberg SL (2012). Fast gapped-read alignment with Bowtie 2. *Nat Meth* **9:** 357-359.

Markowitz VM, Chen IMA, Chu K, Szeto E, Palaniappan K, Grechkin Y *et al* (2012). IMG/M: the integrated metagenome data management and comparative analysis system. *Nucleic Acids Research* **40:** D123-D129.

McDonald D, Price MN, Goodrich J, Nawrocki EP, DeSantis TZ, Probst A *et al* (2012). An improved Greengenes taxonomy with explicit ranks for ecological and evolutionary analyses of bacteria and archaea. *Isme J* **6:** 610-618.

Pruesse E, Quast C, Knittel K, Fuchs BM, Ludwig WG, Peplies J *et al* (2007). SILVA: a comprehensive online resource for quality checked and aligned ribosomal RNA sequence data compatible with ARB. *Nucleic Acids Research* **35:** 7188-7196.

R-Development-Core-Team (2011). R: A language and environment for statistical computing. *R Foundation for Statistical Computing*.

Schloss PD, Westcott SL, Ryabin T, Hall JR, Hartmann M, Hollister EB *et al* (2009). Introducing mothur: Open-Source, Platform-Independent, Community-Supported Software for Describing and Comparing Microbial Communities. *Appl Environ Microb* **75:** 7537-7541.

Takahashi S, Tomita J, Nishioka K, Hisada T, Nishijima M (2014). Development of a prokaryotic universal primer for simultaneous analysis of Bacteria and Archaea using next-generation sequencing. *Plos One* **9:** e105592.

Zhou JZ, Bruns MA, Tiedje JM (1996). DNA recovery from soils of diverse composition. *Appl Environ Microb* **62:** 316-322.

**Supplemental Tables and Figures:**

**Table S1:** Chemical measurements of diffuse fluid from Marker 33 vent in 2015. Percent seawater was calculated via the conservative tracer magnesium (Mg): $[Mg]_{vent}/[Mg]_{seawater}$ *100. Data from Fortunato et al. 2018.

| | **Marker 33 2015** |
|---|---|
| ***Depth (m)*** | 1516 |
| ***Temperature (°C)*** | 40.6 |
| ***Percent Seawater*** | 85.0 |
| ***pH*** | 5.6 |
| ***$H_2S$ (µmol/kg)*** | 627.7 |
| ***$SO_4^{-2}$ (µmol/kg)*** | 22.9 |
| ***$H_2$ (µM)*** | 0.1 |
| ***$CH_4$ (µmol/kg)*** | 27.6 |
| ***$O_2$ (µM)*** | 16.7 |
| ***Microbial cells per mL*** | $7.2 \times 10^5$ |

**Table 2**: Statistics for Epsilonbacteraeota metagenome assembled genomes (MAGs). MAGs were selected manually using tetranucleotide frequency and differential coverage. Percent completeness and contamination was calculated in Anvi'o. For comparison these statistics were also calculated using CheckM.

| **MAG Name** | **Length (Mb)** | **No. of Contigs** | **GC Content** | **ANVI'O Completeness** | **ANVI'O Contamination** | **CheckM Completeness** | **CheckM Contamination** |
|---|---|---|---|---|---|---|---|
| Axial_Epsilon_Bin1 | 0.43 | 289 | 28.7 | 16.1 | 0.0 | 24.5 | 0.0 |
| Axial_Epsilon_Bin2 | 0.37 | 237 | 29.3 | 13.8 | 0.9 | 22.6 | 1.7 |
| Axial_Epsilon_Bin3 | 1.75 | 397 | 47.9 | 93.8 | 4.0 | 96.5 | 2.1 |
| Axial_Epsilon_Bin4 | 0.78 | 226 | 50.2 | 52.0 | 3.3 | 53.7 | 1.7 |
| Axial_Epsilon_Bin5 | 0.62 | 190 | 48.7 | 38.9 | 2.4 | 35.9 | 0.6 |
| Axial_Epsilon_Bin6 | 0.22 | 170 | 47.2 | 18.7 | 1.7 | 20.1 | 0.7 |
| Axial_Epsilon_Bin7 | 0.31 | 179 | 30.9 | 13.3 | 0.0 | 21.2 | 0.0 |
| Axial_Epsilon_Bin8 | 0.27 | 168 | 29.6 | 14.0 | 0.2 | 20.0 | 0.0 |
| Axial_Epsilon_Bin9 | 0.23 | 145 | 28.4 | 22.2 | 0.5 | 21.0 | 0.3 |
| Axial_Epsilon_Bin10 | 1.06 | 367 | 50.1 | 75.2 | 4.1 | 70.2 | 4.3 |

**Table S3:** Results of temperature distribution testing. The experiment was conducted in an ice bath in the lab to determine the steady state temperature distribution throughout the incubation bag relative to the temperature measured at the controlling RTD sensor used in the field deployments. Sensors were placed throughout the bag to measure variation both longitudinally and radially. Longitudinal variation was measured by sensors placed at the controlling RTD sensor, middle and distal end of the bag, and held in place with a custom bracket fabricated to fit snugly within the incubation bag. Radial variation was measured using sensors placed in the longitudinal center, and at the top and bottom surface of the bag. Sensors generally all showed good agreement with each other and with the controlling RTD sensor used to monitor temperature during field deployment. We observed a small but constant offset between bag temperature and controlling RTD temperature, but these lab temperature distribution results are expected to be the worst case, with mixing driven solely by natural convection. In the field experiments there will be additional mixing from vehicle motion. Note, thermocouple numbers reference the RTD temperature probe readings and thermistor numbers represent readings from bag sensors.

| Incubator Setpoint (°C) | Thermocouple Average (°C) | Thermocouple Std. Dev. (°C) | Thermistor Average (°C) | Thermistor Std. Dev. (°C) | Duration (minutes) |
|---|---|---|---|---|---|
| 28 | 28.29 | 0.04 | 30.24 | 0.32 | 10.7 |
| 30 | 30.36 | 0.22 | 32.45 | 0.32 | 17.3 |
| 52 | 50.65 | 0.12 | 53.99 | 0.30 | 5.2 |
| 55 | 55.59 | 0.04 | 58.85 | 0.46 | 5.7 |
| 80 | 80.32 | 0.09 | 84.13 | 0.67 | 7.2 |

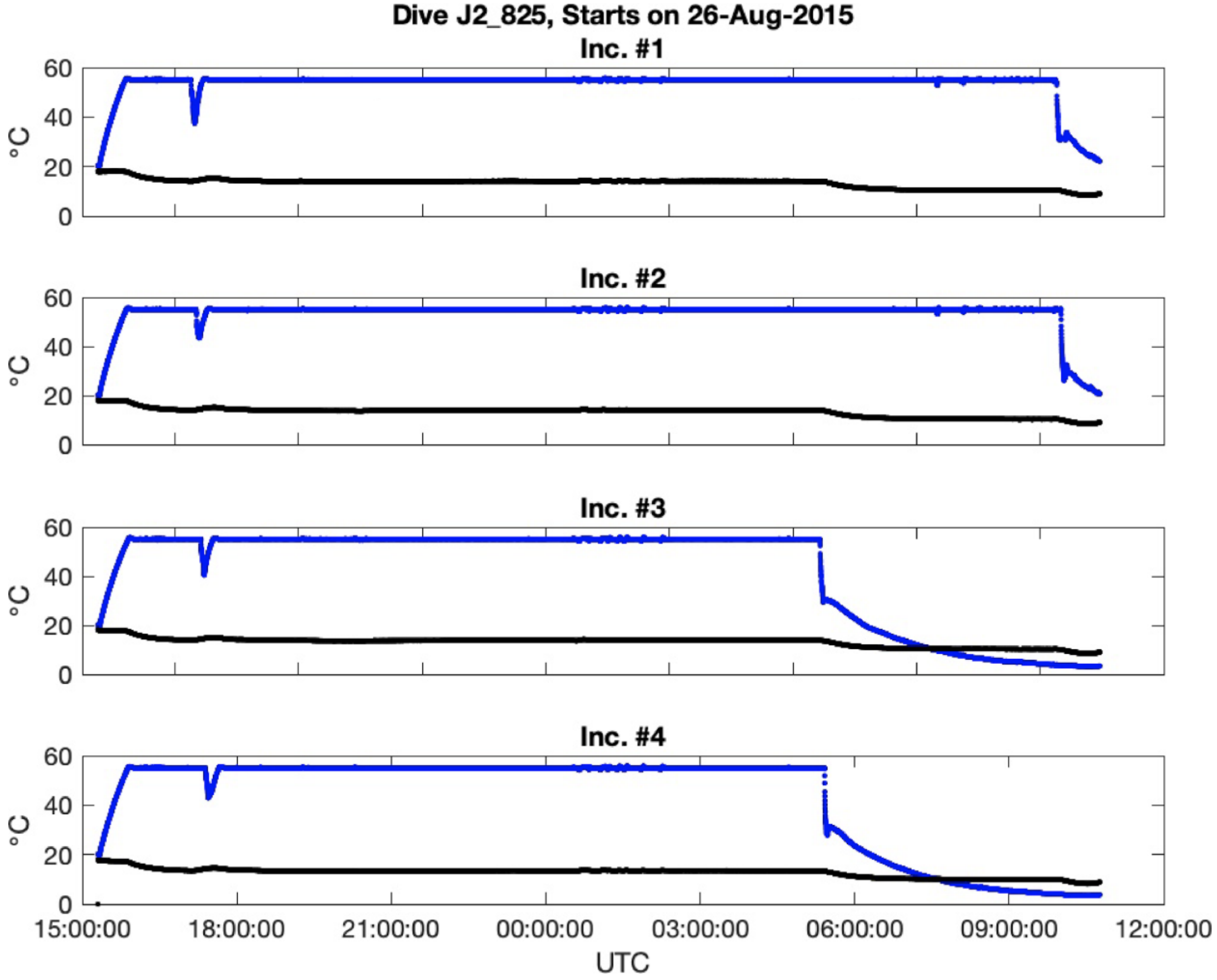


**Figure S1.** Temperature record of the four incubator units during Jason Dive 825 at Marker 33 with all positions heated to 55°C. Blue line shows measured temperature inside the incubation chamber. The brief dip in measured incubation temperature before 18:00 UTC corresponds to intake of 40°C vent fluid into the incubator, marking the beginning of the incubation. Temperatures were very stable throughout the incubation. The black line is the temperature inside the electronics pressure case.

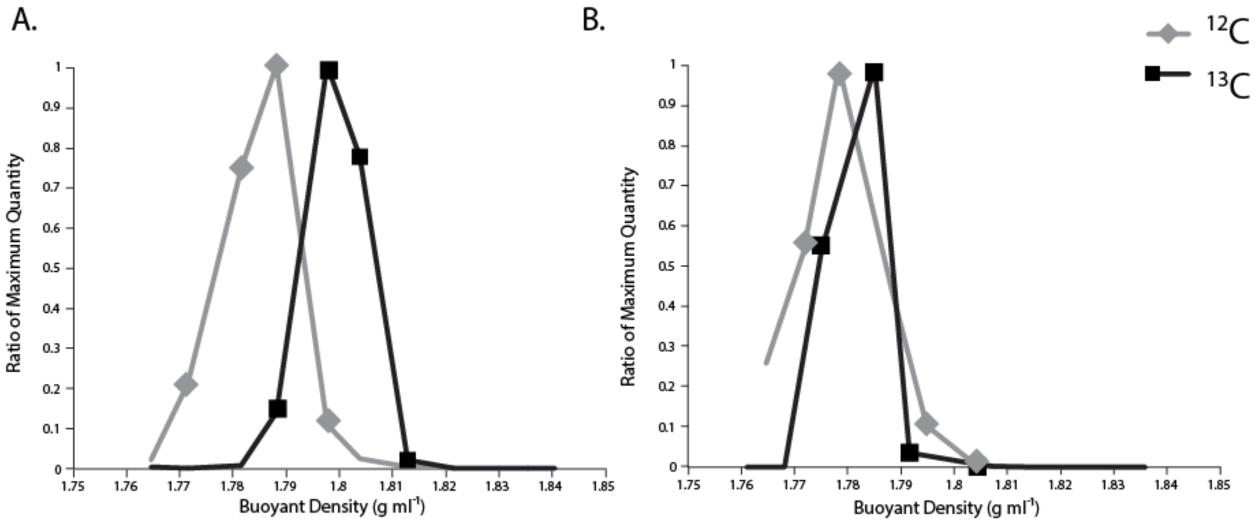


**Figure S2:** Fractionation plots of (A) Shipboard and (B) Incubator RNA-SIP experiments at 16 hours. Buoyant density (g $ml^{-1}$) of each fraction is depicted on the x-axis and amount of 16S rRNA as determined by RT-qPCR is on the y-axis. Amount of 16S rRNA is displayed as the ratio of maximum quantity in order to compare between experiments.

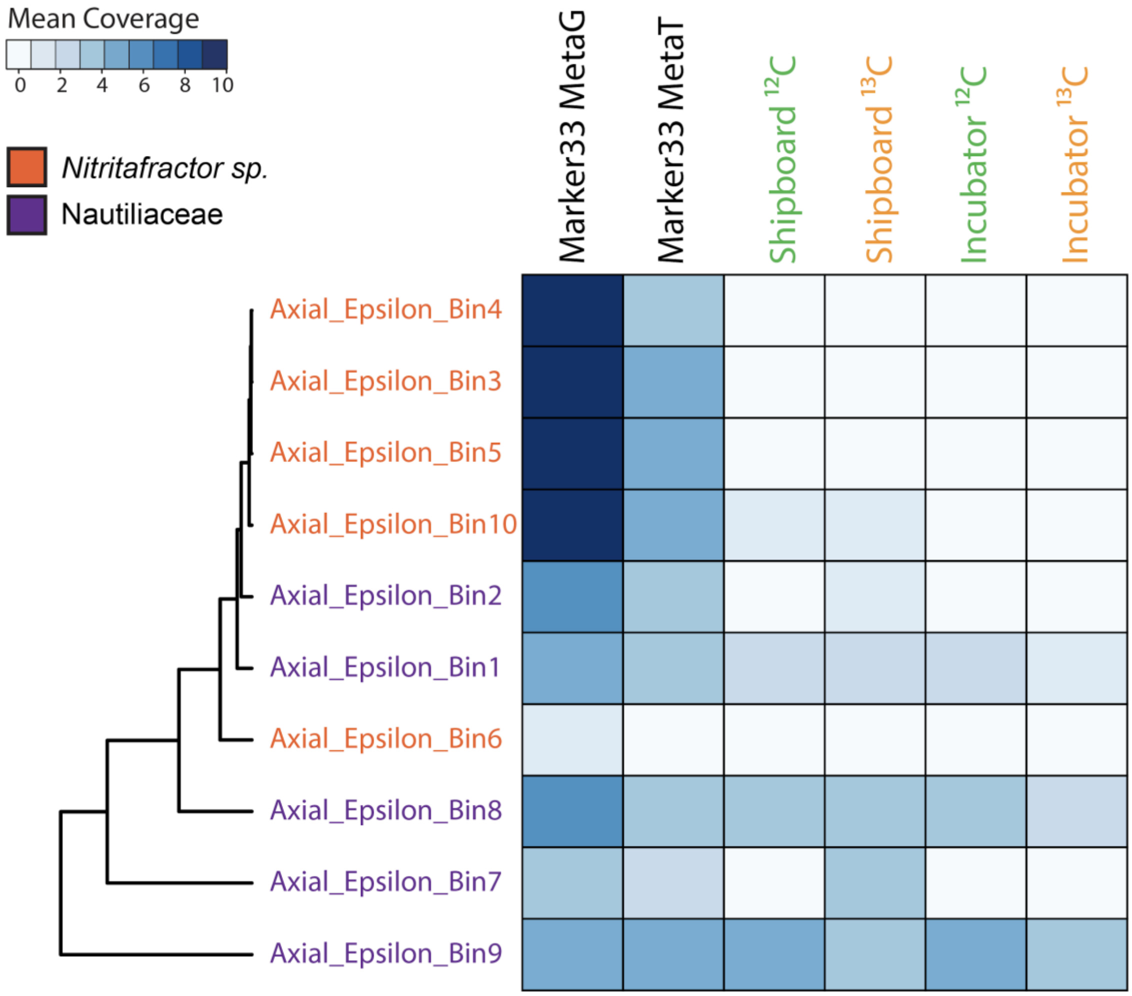


**Figure S3:** Heatmap of mean coverage across the RNA-SIP experiments of metagenome assembled genomes (MAGs) taxonomically identified as thermophilic Epsilonbacteraeota, specifically either the genus *Nitritifactor* (orange) or the family Nautiliaceae (purple). Coverage of bins within the 2015 Marker33 metagenome and metatranscriptome was included for comparison to SIP fractions. Fractions from each of the four RNA-SIP experiments have been collapsed and mean coverage summed. Scale depicts range of mean coverage across MAGs.

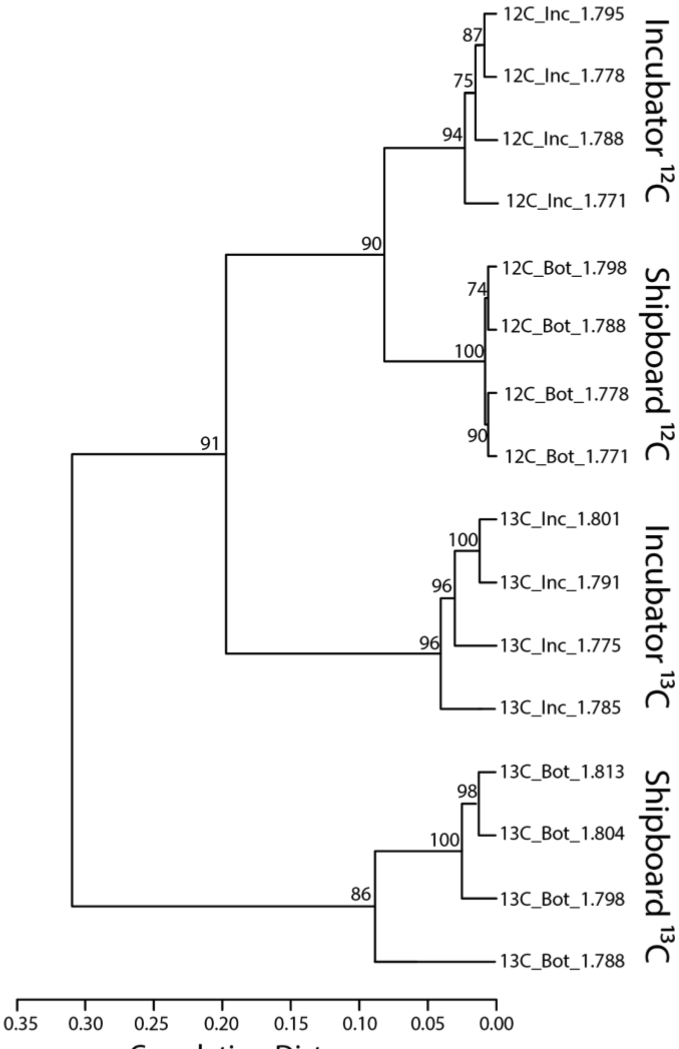


**Figure S4:** Hierarchical clustering of all shipboard and incubator RNA-SIP experiments based on normalized abundance of annotated (KO) transcripts.

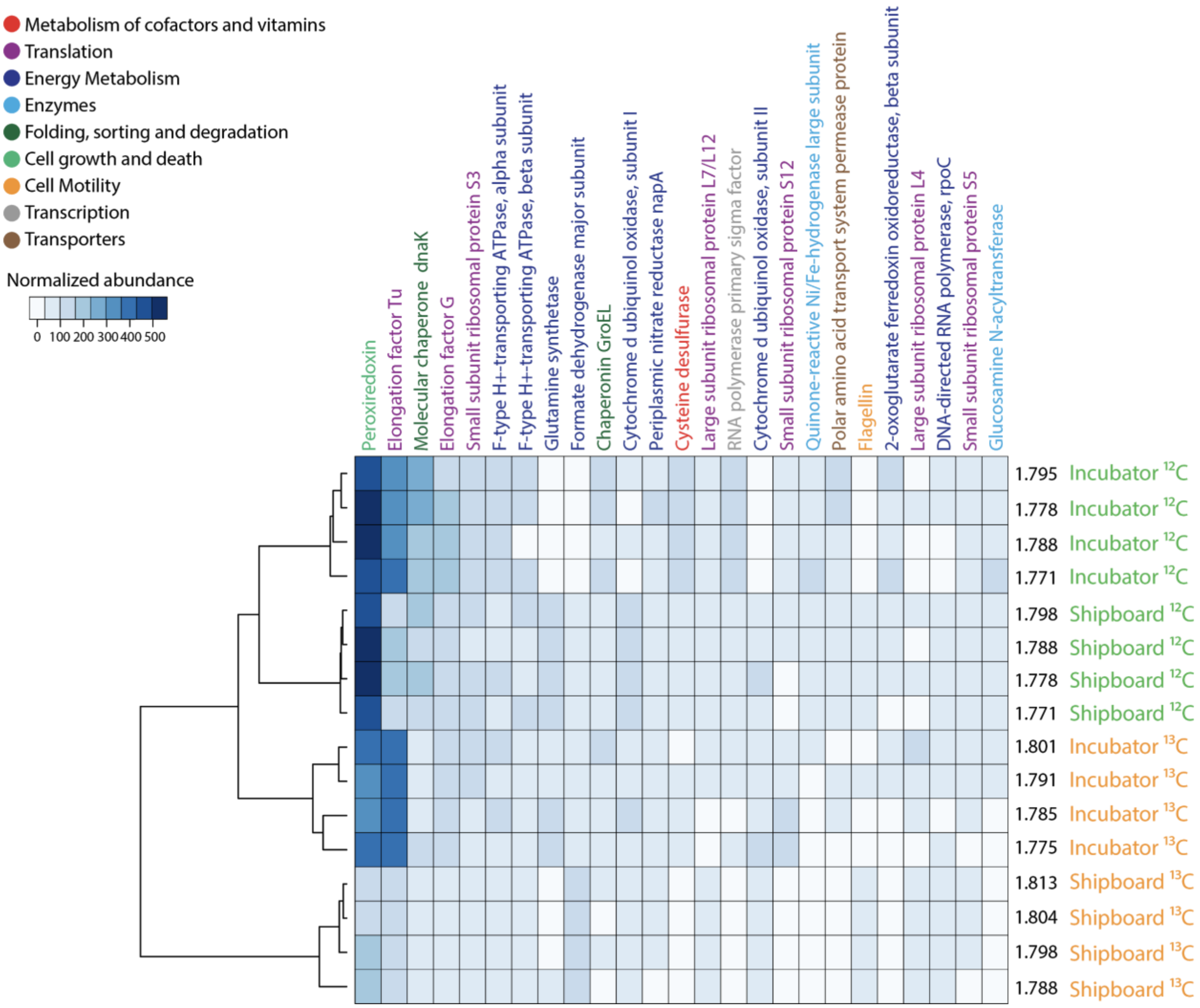


**Figure S5:** Heatmap depicting the twenty-five most abundant annotated transcripts across all RNA-SIP experiments. Transcripts are colored by the KEGG ontology metabolic group. The dendrogram depicts the degree of similarity among the RNA-SIP fractions based on the normalized abundance of the transcripts. Scale depicts the range of normalized abundance, as determined by Transcripts per Million (TPM).

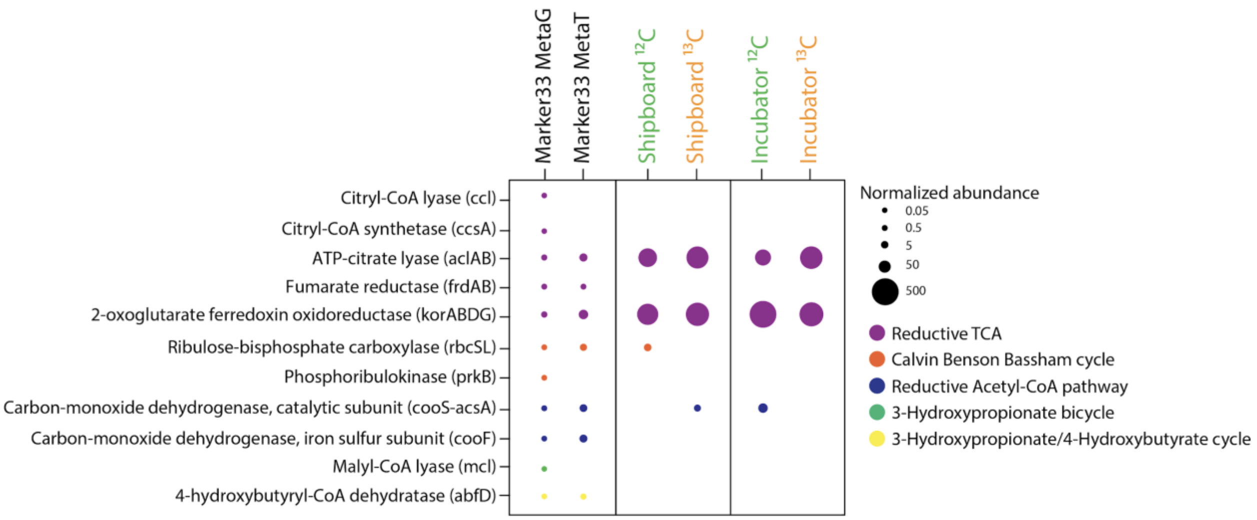


**Figure S6:** Normalized abundance of annotated genes and transcripts for the five main carbon-fixation pathways identified within the 2015 Marker 33 metagenome, metatranscriptome, and the shipboard and seafloor incubator RNA-SIP experiments. Fractions from each of the four RNA-SIP experiments have been collapsed to reflect the normalized abundance of each gene in the entire experiment

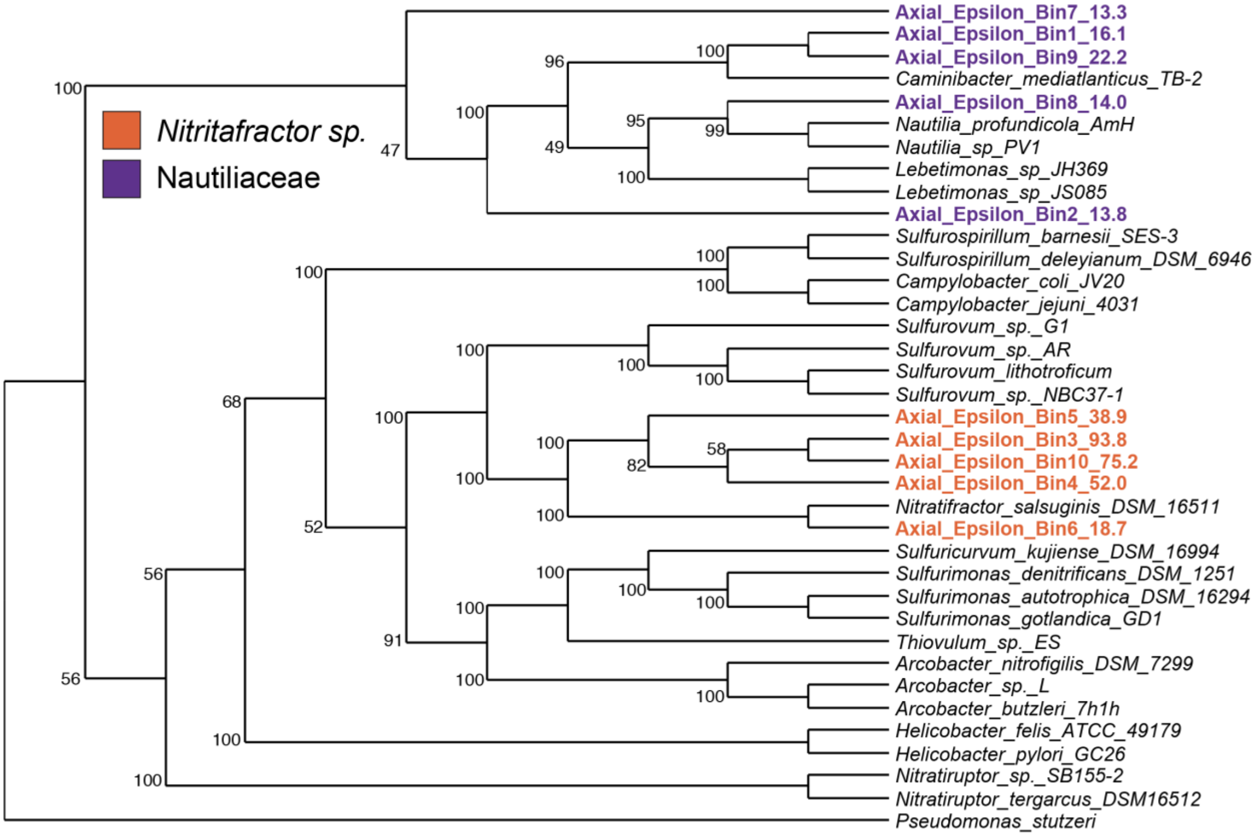


**Figure S7:** Concatenated marker gene tree of thermophilic Epsilonbacteraeota metagenome assembled genomes (MAGs). Amino acid alignments were constructed using the program Phylosift using a reference set of 37 marker genes. Tree was constructed using RAxML. MAG completeness, as determined via CheckM, is denoted at the end of each MAG name. Legend denotes the assigned taxonomy of MAGs as determined in *Anvi'o*.

**Table S4: Significantly differentially expressed transcripts in Shipboard vs. Incubator Experiments (adj p-value < 0.01) as determined by Differential Expression Analysis**

| | | | SHIPBOARD Normalized Abundance | | | | | | | | INCUBATOR Normalized Abundance | | | | | | | | | |
|---|---|---|---|---|---|---|---|---|---|---|---|---|---|---|---|---|---|---|---|---|
| KO_ID | KEGG Gene | KEGG Function | 12C_1.798 | 12C_1.788 | 12C_1.778 | 12C_1.771 | 13C_1.813 | 13C_1.804 | 13C_1.798 | 13C_1.788 | 12C_INC_1.795 | 12C_INC_1.788 | 12C_INC_1.778 | 12C_INC_1.771 | 13C_INC_1.801 | 13C_INC_1.791 | 13C_INC_1.785 | 13C_INC_1.771 | Adjusted p-value | log2FoldChange |
| K03593 | mrp, NUBPL | ATP-binding protein involved in chromosome partitioning | 5691 | 4381 | 3714 | 1856 | 4174 | 3685 | 4852 | 5092 | 0 | 0 | 0 | 0 | 536 | 0 | 0 | 0 | 1.55797E-05 | -11.73066011 |
| K02879 | RP-L17, MRPL17, rplQ | large subunit ribosomal protein L17 | 4752 | 5104 | 655 | 7128 | 4536 | 4627 | 3621 | 2546 | 0 | 0 | 0 | 0 | 3095 | 0 | 0 | 0 | 8.35748E-05 | -11.2088139 |
| K02890 | RP-L22, MRPL22, rplV | large subunit ribosomal protein L22 | 2676 | 2974 | 2450 | 4499 | 3546 | 3669 | 2024 | 2349 | 0 | 0 | 0 | 0 | 1218 | 0 | 0 | 0 | 3.82873E-05 | -11.13282851 |
| K01951 | guaA, GMPS | GMP synthase (glutamine-hydrolysing) [EC:6.3.5.2] | 1876 | 978 | 1840 | 4068 | 2559 | 2260 | 2285 | 2208 | 0 | 0 | 0 | 0 | 697 | 0 | 0 | 0 | 3.89633E-05 | -10.76343784 |
| K02422 | fliS | flagellar protein FliS | 1718 | 1455 | 1573 | 979 | 224 | 518 | 433 | 502 | 0 | 0 | 0 | 0 | 0 | 0 | 0 | 0 | 1.38E-08 | -10.53569074 |
| K03286 | TC.OOP | OmpA-OmpF porin, OOP family | 1163 | 1677 | 4038 | 1991 | 1063 | 1100 | 1193 | 1260 | 0 | 0 | 0 | 0 | 353 | 0 | 0 | 0 | 3.6186E-05 | -10.40517041 |
| K05592 | deaD, cshA | ATP-dependent RNA helicase DeaD [EC:3.6.4.13] | 5159 | 2827 | 4076 | 4593 | 4034 | 4161 | 4545 | 3371 | 0 | 0 | 0 | 0 | 586 | 928 | 0 | 0 | 0.000183526 | -10.3557599 |
| K03455 | TC.KEF | monovalent cation:H+ antiporter-2, CPA2 family | 757 | 1196 | 1180 | 648 | 606 | 507 | 485 | 308 | 0 | 0 | 0 | 0 | 0 | 0 | 0 | 0 | 1.38E-08 | -10.33388812 |
| K01008 | selD, SEPHS | selenide, water dikinase [EC:2.7.9.3] | 2002 | 1569 | 612 | 760 | 2063 | 1764 | 1835 | 1315 | 0 | 0 | 0 | 0 | 293 | 0 | 0 | 0 | 3.6186E-05 | -10.29862445 |
| K02334 | dpo | DNA polymerase bacteriophage-type [EC:2.7.7.7] | 393 | 249 | 720 | 672 | 572 | 650 | 754 | 1179 | 0 | 0 | 0 | 0 | 0 | 0 | 0 | 0 | 1.41E-08 | -10.20336146 |
| K00931 | proB | glutamate 5-kinase [EC:2.7.2.11] | 3089 | 1494 | 1765 | 3172 | 1062 | 836 | 828 | 390 | 0 | 0 | 0 | 0 | 419 | 0 | 0 | 0 | 5.9828E-05 | -10.1399098 |
| K02356 | efp | elongation factor P | 837 | 1329 | 1991 | 716 | 1396 | 1574 | 1233 | 922 | 0 | 0 | 0 | 0 | 303 | 0 | 0 | 0 | 4.61951E-05 | -10.08163796 |
| K06942 | ychF | ribosome-binding ATPase | 431 | 224 | 789 | 737 | 507 | 430 | 844 | 568 | 0 | 0 | 0 | 0 | 0 | 0 | 0 | 0 | 1.41E-08 | -10.02762268 |
| K02357 | tsf, TSFM | elongation factor Ts | 5937 | 5801 | 4918 | 4586 | 3493 | 4213 | 4293 | 2622 | 0 | 0 | 0 | 0 | 1585 | 0 | 0 | 3337 | 0.000486038 | -10.01741794 |
| K08352 | phsA, psrA | thiosulfate reductase / polysulfide reductase chain A | 234 | 736 | 429 | 1204 | 2642 | 2852 | 2514 | 2256 | 0 | 0 | 0 | 0 | 0 | 502 | 0 | 0 | 0.000106787 | -9.968159581 |
| K01933 | purM | phosphoribosylformylglycinamidine cyclo-ligase [EC:6.3.3.1] | 382 | 1033 | 701 | 1310 | 2218 | 2145 | 2323 | 1682 | 0 | 0 | 0 | 0 | 1161 | 0 | 0 | 0 | 0.000175341 | -9.932998372 |
| K02867 | RP-L11, MRPL11, rplK | large subunit ribosomal protein L11 | 581 | 554 | 1065 | 1991 | 1942 | 1889 | 1881 | 1277 | 0 | 0 | 0 | 0 | 1059 | 0 | 0 | 0 | 0.000162141 | -9.929532041 |
| K00076 | hdhA | 7-alpha-hydroxysteroid dehydrogenase [EC:1.1.1.159] | 530 | 1188 | 1512 | 1849 | 1194 | 865 | 1021 | 958 | 0 | 0 | 0 | 0 | 321 | 0 | 0 | 0 | 5.9828E-05 | -9.927224313 |
| K01902 | sucD | succinyl-CoA synthetase alpha subunit [EC:6.2.1.5] | 5883 | 2413 | 898 | 838 | 5790 | 4400 | 5738 | 4663 | 0 | 0 | 0 | 0 | 785 | 1048 | 0 | 0 | 0.000342307 | -9.869544751 |
| K02886 | RP-L2, MRPL2, rplB | large subunit ribosomal protein L2 | 3135 | 2622 | 5741 | 5223 | 3660 | 4194 | 3286 | 1710 | 0 | 0 | 0 | 0 | 634 | 0 | 0 | 3118 | 0.000426416 | -9.867340276 |
| K00145 | argC | N-acetyl-gamma-glutamyl-phosphate reductase [EC:1.2.1.38] | 881 | 956 | 888 | 1020 | 1020 | 869 | 817 | 706 | 0 | 0 | 0 | 0 | 133 | 0 | 0 | 0 | 3.21479E-05 | -9.809304778 |
| K00265 | gltB | glutamate synthase (NADPH/NADH) large chain [EC:1.4.1.13 1. | 2977 | 3630 | 4567 | 4101 | 2385 | 2456 | 2457 | 1947 | 0 | 0 | 0 | 0 | 1026 | 911 | 0 | 0 | 0.000339902 | -9.805409074 |
| K00031 | IDH1, IDH2, icd | isocitrate dehydrogenase [EC:1.1.1.42] | 749 | 2494 | 2632 | 5048 | 6357 | 8403 | 7101 | 5352 | 0 | 0 | 0 | 0 | 2018 | 4381 | 0 | 0 | 0.000816762 | -9.78890176 |
| K00648 | fabH | 3-oxoacyl-[acyl-carrier-protein] synthase III [EC:2.3.1.180] | 1435 | 471 | 1752 | 1801 | 806 | 843 | 720 | 1046 | 0 | 0 | 0 | 0 | 464 | 0 | 0 | 0 | 0.000102058 | -9.77437125 |
| K00171 | porD | pyruvate ferredoxin oxidoreductase delta subunit [EC:1.2.7.1] | 5882 | 3039 | 4370 | 907 | 5531 | 4237 | 3606 | 2624 | 0 | 0 | 0 | 0 | 2927 | 1444 | 0 | 0 | 0.000620499 | -9.715567787 |
| K00818 | E2.6.1.11, argD | acetylornithine aminotransferase [EC:2.6.1.11] | 374 | 1138 | 546 | 641 | 1136 | 1283 | 1165 | 1804 | 0 | 0 | 0 | 0 | 737 | 0 | 0 | 0 | 0.000195395 | -9.53006092 |
| K03590 | ftsA | cell division protein FtsA | 659 | 1226 | 770 | 0 | 1273 | 1137 | 791 | 849 | 0 | 0 | 0 | 0 | 0 | 0 | 0 | 0 | 1.21187E-05 | -9.488773143 |
| K01776 | murI | glutamate racemase [EC:5.1.1.3] | 336 | 320 | 307 | 575 | 687 | 378 | 384 | 147 | 0 | 0 | 0 | 0 | 0 | 0 | 0 | 0 | 3.52E-08 | -9.479854636 |
| K01961 | accC | acetyl-CoA carboxylase, biotin carboxylase subunit [EC:6.4.1.2 6 | 2365 | 2297 | 2950 | 2289 | 1571 | 2180 | 2111 | 1756 | 0 | 0 | 564 | 0 | 1244 | 0 | 0 | 0 | 0.000415948 | -9.414475497 |
| K05927 | hydA3 | quinone-reactive Ni/Fe-hydrogenase small subunit [EC:1.12.5.1 | 1312 | 1367 | 2379 | 648 | 5038 | 5951 | 5951 | 3380 | 0 | 0 | 0 | 0 | 1347 | 2807 | 0 | 0 | 0.001016102 | -9.328529144 |
| K03147 | thiC | phosphomethylpyrimidine synthase [EC:4.1.99.17] | 886 | 1397 | 1826 | 650 | 2780 | 2489 | 2805 | 3322 | 0 | 0 | 0 | 0 | 538 | 474 | 0 | 0 | 0.000362989 | -9.298863427 |
| K13926 | rbbA | ribosome-dependent ATPase | 763 | 445 | 580 | 478 | 645 | 950 | 806 | 598 | 0 | 0 | 0 | 0 | 242 | 0 | 0 | 0 | 0.000115035 | -9.227327212 |
| K03544 | clpX, CLPX | ATP-dependent Clp protease ATP-binding subunit ClpX | 303 | 403 | 464 | 868 | 1024 | 1003 | 1155 | 665 | 0 | 0 | 0 | 0 | 368 | 0 | 0 | 0 | 0.000175341 | -9.212573751 |
| K01867 | WARS, trpS | tryptophanyl-tRNA synthetase [EC:6.1.1.2] | 305 | 534 | 280 | 0 | 1941 | 1748 | 1544 | 537 | 0 | 0 | 0 | 0 | 0 | 0 | 0 | 0 | 1.39816E-05 | -9.208273563 |
| K04771 | degP, htrA | serine protease Do [EC:3.4.21.107] | 380 | 779 | 632 | 1507 | 680 | 627 | 739 | 416 | 0 | 0 | 0 | 0 | 0 | 406 | 0 | 0 | 0.000195395 | -9.173976563 |
| K01677 | E4.2.1.2AA, fumA | fumarate hydratase subunit alpha [EC:4.2.1.2] | 2982 | 1263 | 910 | 0 | 2569 | 2537 | 1622 | 1309 | 0 | 0 | 0 | 0 | 301 | 0 | 0 | 0 | 0.000638944 | -9.155899398 |
| K01265 | map | methionyl aminopeptidase [EC:3.4.11.18] | 568 | 451 | 260 | 486 | 805 | 831 | 750 | 772 | 0 | 0 | 0 | 0 | 173 | 0 | 0 | 0 | 0.000102225 | -9.142272602 |
| K01939 | purA, ADSS | adenylosuccinate synthase [EC:6.3.4.4] | 664 | 658 | 608 | 379 | 500 | 782 | 687 | 681 | 0 | 0 | 0 | 0 | 269 | 0 | 0 | 0 | 0.00014548 | -9.134275674 |
| K13378 | nuoCD | NADH-quinone oxidoreductase subunit C/D [EC:1.6.5.3] | 1183 | 886 | 812 | 0 | 2283 | 2636 | 2352 | 1924 | 0 | 0 | 0 | 0 | 179 | 0 | 0 | 0 | 0.000486038 | -9.117659237 |
| K00341 | nuoL | NADH-quinone oxidoreductase subunit L [EC:1.6.5.3] | 380 | 725 | 488 | 651 | 3687 | 3561 | 2983 | 1952 | 0 | 0 | 0 | 0 | 185 | 326 | 0 | 0 | 0.000339902 | -9.080208195 |
| K00940 | ndk, NME | nucleoside-diphosphate kinase [EC:2.7.4.6] | 0 | 1050 | 585 | 1093 | 501 | 535 | 531 | 561 | 0 | 0 | 0 | 0 | 0 | 0 | 0 | 0 | 1.39816E-05 | -9.052940697 |
| K09010 | K09010 | uncharacterized protein | 2794 | 2471 | 1587 | 1974 | 1030 | 1285 | 1077 | 1073 | 0 | 0 | 0 | 0 | 995 | 409 | 0 | 0 | 0.000542471 | -9.028201639 |
| K01872 | AARS, alaS | alanyl-tRNA synthetase [EC:6.1.1.7] | 1978 | 1543 | 804 | 1928 | 1731 | 1875 | 1928 | 926 | 0 | 0 | 0 | 0 | 961 | 410 | 0 | 0 | 0.000550407 | -8.999184952 |
| K02881 | RP-L18, MRPL18, rplR | large subunit ribosomal protein L18 | 9316 | 11329 | 8563 | 1991 | 4858 | 5401 | 4194 | 2555 | 0 | 1388 | 0 | 0 | 1059 | 2532 | 0 | 0 | 0.001751808 | -8.982844789 |
| K02346 | DPO4, dinB | DNA polymerase IV [EC:2.7.7.7] | 967 | 275 | 590 | 0 | 449 | 506 | 761 | 1123 | 0 | 0 | 0 | 0 | 0 | 0 | 0 | 0 | 1.39816E-05 | -8.949050371 |
| K01736 | aroC | chorismate synthase [EC:4.2.3.5] | 550 | 474 | 0 | 942 | 586 | 728 | 371 | 725 | 0 | 0 | 0 | 0 | 0 | 0 | 0 | 0 | 1.39816E-05 | -8.935302842 |
| K01810 | GPI, pgi | glucose-6-phosphate isomerase [EC:5.3.1.9] | 315 | 556 | 606 | 532 | 670 | 629 | 614 | 138 | 0 | 0 | 0 | 0 | 104 | 0 | 0 | 0 | 9.23745E-05 | -8.90517473 |
| K01678 | E4.2.1.2AB, fumB | fumarate hydratase subunit beta [EC:4.2.1.2] | 773 | 1351 | 2834 | 0 | 2202 | 2196 | 1931 | 1189 | 0 | 0 | 0 | 0 | 939 | 0 | 0 | 0 | 0.001574803 | -8.884004478 |
| K00241 | sdhC, frdC | succinate dehydrogenase / fumarate reductase, cytochrome b | 1620 | 1819 | 216 | 1616 | 2893 | 3103 | 2458 | 2570 | 0 | 0 | 0 | 0 | 997 | 0 | 0 | 1410 | 0.001341925 | -8.865557024 |
| K02952 | RP-S13, rpsM | small subunit ribosomal protein S13 | 4078 | 2495 | 4263 | 4979 | 3028 | 4569 | 3786 | 5366 | 0 | 0 | 510 | 0 | 709 | 2500 | 0 | 0 | 0.001386598 | -8.856803148 |
| K03770 | ppiD | peptidyl-prolyl cis-trans isomerase D [EC:5.2.1.8] | 666 | 315 | 852 | 451 | 514 | 652 | 775 | 599 | 0 | 0 | 0 | 0 | 885 | 0 | 0 | 0 | 0.000362989 | -8.850065565 |
| K02933 | RP-L6, MRPL6, rplF | large subunit ribosomal protein L6 | 7465 | 6657 | 5550 | 5631 | 4498 | 3806 | 3554 | 1436 | 0 | 0 | 0 | 0 | 564 | 1014 | 4221 | 0 | 0.001962719 | -8.816283096 |
| K09861 | K09861 | uncharacterized protein | 614 | 341 | 281 | 0 | 624 | 545 | 712 | 874 | 0 | 0 | 0 | 0 | 0 | 0 | 0 | 0 | 1.47646E-05 | -8.787440062 |
| K02878 | RP-L16, MRPL16, rplP | large subunit ribosomal protein L16 | 1120 | 1359 | 537 | 0 | 1676 | 2244 | 1659 | 984 | 0 | 0 | 0 | 0 | 356 | 0 | 0 | 0 | 0.001062074 | -8.721110945 |
| K00615 | E2.2.1.1, tktA, tktB | transketolase [EC:2.2.1.1] | 1573 | 1233 | 1074 | 2052 | 2172 | 1696 | 1281 | 930 | 0 | 0 | 0 | 0 | 1044 | 1516 | 0 | 0 | 0.001379935 | -8.68008002 |
| K02495 | hemN, hemZ | oxygen-independent coproporphyrinogen III oxidase [EC:1.3.99. | 1502 | 788 | 708 | 0 | 1368 | 1689 | 1595 | 761 | 0 | 0 | 0 | 0 | 234 | 0 | 0 | 0 | 0.000868666 | -8.659186215 |
| K03924 | moxR | MoxR-like ATPase [EC:3.6.3.-] | 897 | 350 | 1896 | 2058 | 943 | 670 | 807 | 1001 | 0 | 0 | 122 | 0 | 0 | 589 | 0 | 0 | 0.000486038 | -8.627277745 |
| K03687 | GRPE | molecular chaperone GrpE | 1645 | 1306 | 753 | 703 | 1776 | 2435 | 2240 | 1264 | 0 | 0 | 0 | 0 | 1248 | 1760 | 0 | 0 | 0.001762203 | -8.565397392 |
| K00124 | fdoH | formate dehydrogenase iron-sulfur subunit | 4676 | 4570 | 5600 | 2462 | 7202 | 8303 | 5754 | 4414 | 0 | 0 | 0 | 0 | 3493 | 2308 | 0 | 4188 | 0.002991887 | -8.550155491 |
| K01092 | E3.1.3.25, IMPA, suhB | myo-inositol-1(or 4)-monophosphatase [EC:3.1.3.25] | 627 | 903 | 1060 | 0 | 1447 | 1571 | 1171 | 882 | 0 | 0 | 0 | 0 | 257 | 0 | 0 | 0 | 0.001007112 | -8.549948012 |
| K00919 | ispE | 4-diphosphocytidyl-2-C-methyl-D-erythritol kinase [EC:2.7.1.14 | 1392 | 800 | 642 | 0 | 140 | 300 | 520 | 208 | 0 | 0 | 0 | 0 | 0 | 0 | 0 | 0 | 1.55797E-05 | -8.549769129 |
| K01749 | hemC, HMBS | hydroxymethylbilane synthase [EC:2.5.1.61] | 3113 | 1126 | 516 | 1484 | 2093 | 1431 | 1174 | 1156 | 0 | 0 | 0 | 0 | 457 | 0 | 3939 | 0 | 0.001962719 | -8.52880742 |
| K06207 | typA, bipA | GTP-binding protein | 2719 | 2210 | 2575 | 682 | 1429 | 1920 | 1544 | 991 | 0 | 0 | 0 | 0 | 1987 | 0 | 0 | 4484 | 0.002404388 | -8.486101313 |
| K02004 | ABC.CD.P | putative ABC transport system permease protein | 0 | 337 | 697 | 382 | 629 | 541 | 443 | 135 | 0 | 0 | 0 | 0 | 0 | 0 | 0 | 0 | 1.55797E-05 | -8.445250716 |
| K03587 | ftsI | cell division protein FtsI (penicillin-binding protein 3) | 772 | 327 | 235 | 0 | 462 | 419 | 344 | 443 | 0 | 0 | 0 | 0 | 0 | 0 | 0 | 0 | 1.55797E-05 | -8.435459599 |
| K00645 | fabD | [acyl-carrier-protein] S-malonyltransferase [EC:2.3.1.39] | 376 | 87 | 345 | 0 | 429 | 601 | 772 | 668 | 0 | 0 | 0 | 0 | 0 | 0 | 0 | 0 | 1.55797E-05 | -8.406594637 |
| K10857 | exoX | exodeoxyribonuclease X [EC:3.1.11.-] | 798 | 1268 | 329 | 1366 | 121 | 190 | 126 | 0 | 0 | 0 | 0 | 0 | 0 | 0 | 0 | 0 | 1.55797E-05 | -8.400121106 |
| K00557 | trmA | tRNA (uracil-5-)-methyltransferase [EC:2.1.1.35] | 837 | 191 | 743 | 380 | 217 | 215 | 340 | 172 | 0 | 0 | 0 | 0 | 119 | 0 | 0 | 0 | 0.000212357 | -8.378268501 |
| K00795 | ispA | farnesyl diphosphate synthase [EC:2.5.1.1 2.5.1.10] | 847 | 201 | 775 | 0 | 207 | 282 | 289 | 557 | 0 | 0 | 0 | 0 | 0 | 0 | 0 | 0 | 1.55797E-05 | -8.374120112 |
| K00290 | LYS1 | saccharopine dehydrogenase (NAD+, L-lysine forming) [EC:1.5. | 493 | 1004 | 818 | 0 | 902 | 941 | 1360 | 1278 | 0 | 0 | 0 | 0 | 291 | 0 | 0 | 0 | 0.001319295 | -8.372512015 |
| K03601 | xseA | exodeoxyribonuclease VII large subunit [EC:3.1.11.6] | 795 | 93 | 449 | 453 | 261 | 359 | 446 | 490 | 0 | 0 | 0 | 0 | 0 | 343 | 0 | 0 | 0.000392953 | -8.340731436 |
| K02406 | fliC | flagellin | 6545 | 8211 | 10191 | 11298 | 11325 | 12480 | 13613 | 10475 | 0 | 3387 | 705 | 0 | 3594 | 2090 | 0 | 0 | 0.002991887 | -8.31577392 |
| K03100 | lepB | signal peptidase I [EC:3.4.21.89] | 872 | 207 | 399 | 746 | 128 | 302 | 363 | 0 | 0 | 0 | 0 | 0 | 0 | 0 | 0 | 0 | 1.55797E-05 | -8.297142615 |
| K03798 | ftsH, hflB | cell division protease FtsH [EC:3.4.24.-] | 668 | 325 | 1306 | 0 | 2556 | 2383 | 1640 | 1576 | 0 | 0 | 0 | 0 | 0 | 0 | 0 | 2044 | 0.003245998 | -8.276223064 |
| K01963 | accD | acetyl-CoA carboxylase carboxyl transferase subunit beta [EC:6 | 1553 | 576 | 0 | 703 | 859 | 1205 | 1135 | 361 | 0 | 0 | 0 | 0 | 249 | 0 | 0 | 0 | 0.001342239 | -8.256098559 |
| K00127 | fdoI | formate dehydrogenase subunit gamma | 3925 | 2640 | 2537 | 1580 | 7978 | 8807 | 7265 | 3852 | 0 | 0 | 0 | 0 | 1822 | 2964 | 6290 | 0 | 0.004131099 | -8.243537969 |
| K03750 | moeA | molybdopterin molybdotransferase [EC:2.10.1.1] | 1052 | 731 | 959 | 808 | 1680 | 1482 | 1073 | 970 | 0 | 0 | 0 | 0 | 663 | 0 | 2037 | 0 | 0.002057842 | -8.229928863 |
| K02897 | RP-L25, rplY | large subunit ribosomal protein L25 | 3812 | 2354 | 4266 | 3623 | 5069 | 2977 | 4858 | 3363 | 0 | 0 | 0 | 0 | 3815 | 906 | 4887 | 0 | 0.003604719 | -8.214307875 |
| K05808 | yhbH | putative sigma-54 modulation protein | 428 | 680 | 785 | 0 | 883 | 1427 | 1296 | 1129 | 0 | 0 | 0 | 0 | 520 | 0 | 0 | 0 | 0.001996857 | -8.206932159 |
| K02453 | gspD | general secretion pathway protein D | 511 | 270 | 1560 | 583 | 801 | 1033 | 1009 | 903 | 0 | 0 | 0 | 0 | 310 | 364 | 0 | 0 | 0.000940922 | -8.201712372 |
| K09760 | rmuC | DNA recombination protein RmuC | 493 | 206 | 512 | 0 | 248 | 471 | 233 | 325 | 0 | 0 | 0 | 0 | 0 | 0 | 0 | 0 | 1.55797E-05 | -8.195827547 |
| K06168 | miaB | tRNA-2-methylthio-N6-dimethylallyladenosine synthase [EC:2. | 618 | 405 | 986 | 760 | 948 | 837 | 729 | 640 | 0 | 0 | 0 | 0 | 269 | 372 | 0 | 0 | 0.000882299 | -8.168402383 |
| K03408 | cheW | purine-binding chemotaxis protein CheW | 0 | 462 | 762 | 712 | 815 | 1004 | 1099 | 727 | 0 | 0 | 0 | 0 | 251 | 0 | 0 | 0 | 0.001386598 | -8.16619598 |
| K03620 | hyaC | Ni/Fe-hydrogenase 1 B-type cytochrome subunit | 370 | 176 | 0 | 634 | 1958 | 1190 | 1032 | 1465 | 0 | 0 | 0 | 0 | 225 | 0 | 0 | 0 | 0.00145926 | -8.161667611 |
| K02528 | ksgA | 16S rRNA (adenine1518-N6/adenine1519-N6)-dimethyltransfe | 1645 | 172 | 376 | 703 | 946 | 774 | 1101 | 2102 | 0 | 0 | 0 | 0 | 219 | 880 | 0 | 0 | 0.001386598 | -8.156288093 |
| K06039 | ychN | uncharacterized protein involved in oxidation of intracellular su | 1203 | 191 | 550 | 0 | 1298 | 1111 | 1137 | 792 | 0 | 0 | 0 | 0 | 365 | 0 | 0 | 0 | 0.001961308 | -8.099484933 |

| | | | | | | | | | | | | | | | | | | | | |
|---|---|---|---|---|---|---|---|---|---|---|---|---|---|---|---|---|---|---|---|---|
| K01610 | E4.1.1.49, pckA | phosphoenolpyruvate carboxykinase (ATP) [EC:4.1.1.49] | 3656 | 4068 | 1947 | 641 | 3695 | 4425 | 4126 | 4406 | 0 | 0 | 0 | 0 | 1475 | 801 | 0 | 4473 | 0.003500785 | -8.089527403 |
| K02996 | RP-S9, MRPS9, rpsI | small subunit ribosomal protein S9 | 0 | 766 | 736 | 1376 | 552 | 588 | 1136 | 706 | 0 | 0 | 0 | 0 | 488 | 0 | 0 | 0 | 0.002044762 | -8.087080602 |
| K01687 | ilvD | dihydroxy-acid dehydratase [EC:4.2.1.9] | 2257 | 1002 | 118 | 811 | 1389 | 1773 | 2130 | 1390 | 0 | 0 | 0 | 0 | 852 | 0 | 4305 | 0 | 0.002991887 | -8.052375639 |
| K00172 | porG | pyruvate ferredoxin oxidoreductase gamma subunit [EC:1.2.7.1 | 586 | 652 | 2149 | 0 | 863 | 711 | 621 | 772 | 0 | 0 | 0 | 0 | 712 | 0 | 0 | 0 | 0.002331252 | -8.049497824 |
| K08303 | K08303 | putative protease [EC:3.4.-.-] | 1722 | 597 | 0 | 2401 | 594 | 672 | 451 | 758 | 0 | 0 | 0 | 0 | 996 | 0 | 0 | 0 | 0.002498768 | -8.048109395 |
| K02874 | RP-L14, MRPL14, rplN | large subunit ribosomal protein L14 | 581 | 3485 | 3730 | 2987 | 2056 | 2787 | 1899 | 2044 | 0 | 1377 | 519 | 0 | 983 | 0 | 0 | 0 | 0.002475814 | -8.039929773 |
| K01433 | purU | formyltetrahydrofolate deformylase [EC:3.5.1.10] | 431 | 479 | 394 | 0 | 216 | 178 | 130 | 378 | 0 | 0 | 0 | 0 | 0 | 0 | 0 | 0 | 1.98384E-05 | -7.999812153 |
| K01698 | hemB, ALAD | porphobilinogen synthase [EC:4.2.1.24] | 461 | 292 | 0 | 1359 | 1119 | 1144 | 1000 | 998 | 0 | 0 | 0 | 0 | 898 | 0 | 0 | 0 | 0.002498768 | -7.998863599 |
| K00286 | proC | pyrroline-5-carboxylate reductase [EC:1.5.1.2] | 290 | 369 | 1165 | 0 | 651 | 840 | 795 | 822 | 0 | 0 | 0 | 0 | 176 | 0 | 0 | 0 | 0.001386598 | -7.993573778 |
| K01696 | trpB | tryptophan synthase beta chain [EC:4.2.1.20] | 680 | 673 | 902 | 1123 | 1323 | 739 | 656 | 731 | 0 | 0 | 0 | 0 | 335 | 0 | 2509 | 0 | 0.002331252 | -7.960618429 |
| K18682 | rny | ribonucrease Y [EC:3.1.-.-] | 144 | 251 | 395 | 492 | 718 | 989 | 957 | 1151 | 0 | 0 | 0 | 0 | 98 | 308 | 0 | 0 | 0.000878281 | -7.924604638 |
| K00088 | guaB | IMP dehydrogenase [EC:1.1.1.205] | 299 | 1267 | 659 | 512 | 658 | 969 | 917 | 924 | 0 | 0 | 0 | 0 | 437 | 900 | 0 | 0 | 0.001996857 | -7.919405053 |
| K08602 | pepF, pepB | oligoendopeptidase F [EC:3.4.24.-] | 0 | 100 | 193 | 362 | 428 | 439 | 643 | 92 | 0 | 0 | 0 | 0 | 0 | 0 | 0 | 0 | 2.50022E-05 | -7.877716924 |
| K12573 | rnr, vacB | ribonuclease R [EC:3.1.-.-] | 483 | 422 | 1036 | 0 | 331 | 730 | 563 | 601 | 0 | 0 | 0 | 0 | 146 | 0 | 0 | 0 | 0.001386598 | -7.856357061 |
| K00820 | glmS, GFPT | glucosamine--fructose-6-phosphate aminotransferase (isomer | 0 | 409 | 822 | 1480 | 862 | 720 | 400 | 302 | 0 | 0 | 0 | 0 | 325 | 0 | 0 | 0 | 0.002049571 | -7.855479728 |
| K00169 | porA | pyruvate ferredoxin oxidoreductase alpha subunit [EC:1.2.7.1] | 3526 | 4044 | 5651 | 2715 | 9011 | 9312 | 7986 | 7137 | 0 | 415 | 0 | 0 | 2514 | 1131 | 1601 | 0 | 0.003443856 | -7.854168909 |
| K02600 | nusA | N utilization substance protein A | 708 | 292 | 321 | 0 | 971 | 745 | 890 | 1346 | 0 | 0 | 0 | 0 | 529 | 0 | 0 | 0 | 0.002331252 | -7.843939809 |
| K01924 | murC | UDP-N-acetylmuramate--alanine ligase [EC:6.3.2.8] | 522 | 386 | 239 | 894 | 1344 | 1326 | 1037 | 1138 | 0 | 0 | 0 | 0 | 458 | 0 | 1497 | 0 | 0.002371953 | -7.839366951 |
| K06178 | rluB | 23S rRNA pseudouridine2605 synthase [EC:5.4.99.22] | 774 | 614 | 0 | 624 | 572 | 528 | 360 | 1111 | 0 | 0 | 0 | 0 | 374 | 0 | 0 | 0 | 0.002163997 | -7.820440981 |
| K06180 | rluD | 23S rRNA pseudouridine1911/1915/1917 synthase [EC:5.4.99.2 | 0 | 465 | 402 | 0 | 1049 | 602 | 1006 | 894 | 0 | 0 | 0 | 0 | 0 | 0 | 0 | 0 | 0.00021558 | -7.805082236 |
| K02481 | K02481 | two-component system, NtrC family, response regulator | 2324 | 664 | 1277 | 2386 | 329 | 300 | 267 | 0 | 0 | 0 | 0 | 0 | 0 | 1486 | 0 | 0 | 0.003389274 | -7.798823965 |
| K00759 | APRT, apt | adenine phosphoribosyltransferase [EC:2.4.2.7] | 406 | 452 | 0 | 695 | 915 | 688 | 676 | 357 | 0 | 0 | 0 | 0 | 246 | 0 | 0 | 0 | 0.001969265 | -7.784775133 |
| K01998 | livM | branched-chain amino acid transport system permease proteir | 1045 | 417 | 0 | 346 | 811 | 900 | 913 | 370 | 0 | 0 | 0 | 0 | 493 | 0 | 0 | 0 | 0.002331252 | -7.779335224 |
| K01937 | pyrG, CTPS | CTP synthase [EC:6.3.4.2] | 740 | 938 | 693 | 0 | 881 | 583 | 751 | 166 | 0 | 0 | 0 | 0 | 443 | 0 | 0 | 0 | 0.002331252 | -7.766883111 |
| K00003 | E1.1.1.3 | homoserine dehydrogenase [EC:1.1.1.3] | 550 | 206 | 282 | 0 | 589 | 597 | 761 | 1560 | 0 | 0 | 0 | 0 | 187 | 0 | 0 | 0 | 0.001847139 | -7.763847384 |
| K02470 | gyrB | DNA gyrase subunit B [EC:5.99.1.3] | 1187 | 2097 | 2505 | 1331 | 2083 | 1889 | 1582 | 2143 | 0 | 0 | 0 | 0 | 425 | 1086 | 0 | 1509 | 0.003028809 | -7.750932245 |
| K03111 | ssb | single-strand DNA-binding protein | 3171 | 1942 | 3734 | 775 | 3845 | 3491 | 3495 | 5031 | 0 | 0 | 0 | 0 | 2562 | 2907 | 4114 | 0 | 0.007003935 | -7.727627836 |
| K00791 | miaA, TRIT1 | tRNA dimethylallyltransferase [EC:2.5.1.75] | 316 | 417 | 310 | 580 | 66 | 130 | 152 | 0 | 0 | 0 | 0 | 0 | 0 | 0 | 0 | 0 | 2.40865E-05 | -7.714832088 |
| K07090 | K07090 | uncharacterized protein | 0 | 372 | 546 | 0 | 986 | 522 | 739 | 736 | 0 | 0 | 0 | 0 | 0 | 0 | 0 | 0 | 0.000224617 | -7.694181497 |
| K03076 | secY | preprotein translocase subunit SecY | 4362 | 3478 | 2810 | 2936 | 5341 | 4199 | 3154 | 3469 | 0 | 402 | 0 | 1081 | 238 | 0 | 1444 | 0 | 0.00288115 | -7.675097983 |
| K02563 | murG | UDP-N-acetylglucosamine--N-acetylmuramyl-(pentapeptide) p | 483 | 71 | 196 | 0 | 307 | 247 | 201 | 208 | 0 | 0 | 0 | 0 | 0 | 0 | 0 | 0 | 2.76459E-05 | -7.645828435 |
| K02109 | ATPF0B, atpF | F-type H+-transporting ATPase subunit b | 423 | 605 | 1268 | 2174 | 899 | 1215 | 780 | 472 | 0 | 0 | 0 | 0 | 820 | 0 | 0 | 7584 | 0.004955875 | -7.636619501 |
| K04567 | KARS, lysS | lysyl-tRNA synthetase, class II [EC:6.1.1.6] | 196 | 404 | 718 | 335 | 907 | 962 | 856 | 1624 | 0 | 0 | 0 | 0 | 357 | 1799 | 0 | 0 | 0.002643461 | -7.632450357 |
| K15495 | wtpA | molybdate/tungstate transport system substrate-binding prote | 365 | 911 | 1024 | 0 | 1676 | 2557 | 2408 | 1964 | 0 | 0 | 185 | 0 | 561 | 0 | 0 | 0 | 0.004760707 | -7.623552339 |
| K02388 | flgC | flagellar basal-body rod protein FlgC | 688 | 437 | 0 | 0 | 804 | 366 | 714 | 604 | 0 | 0 | 0 | 0 | 0 | 0 | 0 | 0 | 0.000227235 | -7.623022072 |
| K11392 | rsmF | 16S rRNA (cytosine1407-C5)-methyltransferase [EC:2.1.1.178] | 490 | 456 | 0 | 0 | 615 | 414 | 585 | 1064 | 0 | 0 | 0 | 0 | 0 | 0 | 0 | 0 | 0.000228866 | -7.609805768 |
| K03582 | recB | exodeoxyribonuclease V beta subunit [EC:3.1.11.5] | 189 | 127 | 0 | 324 | 147 | 184 | 141 | 568 | 0 | 0 | 0 | 0 | 0 | 0 | 0 | 0 | 2.82647E-05 | -7.603663528 |
| K01754 | E4.3.1.19, ilvA, tdcB | threonine dehydratase [EC:4.3.1.19] | 511 | 475 | 345 | 1029 | 536 | 785 | 744 | 770 | 0 | 0 | 0 | 0 | 916 | 808 | 0 | 0 | 0.002611491 | -7.589443775 |
| K02519 | infB, MTIF2 | translation initiation factor IF-2 | 483 | 1186 | 1293 | 760 | 1681 | 2113 | 2336 | 1730 | 0 | 0 | 75 | 0 | 1117 | 1698 | 0 | 0 | 0.002991887 | -7.580682473 |
| K02015 | ABC.FEV.P | iron complex transport system permease protein | 218 | 138 | 199 | 0 | 299 | 184 | 66 | 670 | 0 | 0 | 0 | 0 | 0 | 0 | 0 | 0 | 3.11797E-05 | -7.578255903 |
| K03070 | secA | preprotein translocase subunit SecA | 1641 | 1447 | 0 | 695 | 1392 | 1633 | 1677 | 1525 | 0 | 386 | 0 | 0 | 498 | 0 | 0 | 0 | 0.005792399 | -7.565527362 |
| K00928 | lysC | aspartate kinase [EC:2.7.2.4] | 596 | 450 | 546 | 1020 | 690 | 619 | 670 | 931 | 0 | 0 | 0 | 0 | 741 | 0 | 0 | 1865 | 0.003248763 | -7.554427078 |
| K07576 | K07576 | metallo-beta-lactamase family protein | 1093 | 288 | 0 | 594 | 519 | 652 | 707 | 661 | 0 | 0 | 0 | 0 | 1367 | 0 | 0 | 0 | 0.003875128 | -7.551899389 |
| K02469 | gyrA | DNA gyrase subunit A [EC:5.99.1.3] | 1327 | 1453 | 1023 | 964 | 1133 | 1194 | 1233 | 1138 | 0 | 0 | 129 | 770 | 760 | 0 | 0 | 0 | 0.002504319 | -7.54708577 |
| K00981 | E2.7.7.41, CDS1, CDS2, cd | phosphatidate cytidylyltransferase [EC:2.7.7.41] | 530 | 279 | 485 | 0 | 718 | 404 | 383 | 465 | 0 | 0 | 0 | 0 | 176 | 0 | 0 | 0 | 0.001996857 | -7.546073227 |
| K02433 | gatA, QRSL1 | aspartyl-tRNA(Asn)/glutamyl-tRNA(Gln) amidotransferase sut | 365 | 319 | 668 | 0 | 615 | 586 | 509 | 160 | 0 | 0 | 0 | 0 | 110 | 0 | 0 | 0 | 0.001583442 | -7.543064387 |
| K11755 | hisIE | phosphoribosyl-ATP pyrophosphohydrolase / phosphoribosyl-AI | 1001 | 78 | 455 | 0 | 469 | 621 | 781 | 623 | 0 | 0 | 0 | 0 | 258 | 0 | 0 | 0 | 0.002331252 | -7.533275921 |
| K02397 | flgL | flagellar hook-associated protein 3 FlgL | 0 | 239 | 187 | 349 | 956 | 657 | 678 | 723 | 0 | 0 | 0 | 0 | 249 | 0 | 0 | 0 | 0.002312648 | -7.530030917 |
| K00616 | E2.2.1.2, talA, talB | transaldolase [EC:2.2.1.2] | 0 | 285 | 929 | 0 | 388 | 521 | 627 | 651 | 0 | 0 | 0 | 0 | 0 | 0 | 0 | 0 | 0.000237975 | -7.52403111 |
| K03551 | ruvB | holliday junction DNA helicase RuvB [EC:3.6.4.12] | 313 | 498 | 575 | 0 | 1130 | 965 | 748 | 827 | 0 | 0 | 0 | 0 | 0 | 0 | 0 | 1873 | 0.005867488 | -7.51396985 |
| K07024 | K07024 | uncharacterized protein | 264 | 293 | 241 | 451 | 94 | 101 | 119 | 0 | 0 | 0 | 0 | 0 | 0 | 0 | 0 | 0 | 2.64593E-05 | -7.499498264 |
| K09815 | znuA | zinc transport system substrate-binding protein | 477 | 347 | 0 | 935 | 563 | 519 | 670 | 659 | 0 | 0 | 0 | 0 | 0 | 1169 | 0 | 0 | 0.003875128 | -7.495232503 |
| K06196 | ccdA | cytochrome c-type biogenesis protein | 804 | 383 | 0 | 0 | 959 | 425 | 779 | 176 | 0 | 0 | 0 | 0 | 0 | 0 | 0 | 0 | 0.000245274 | -7.481133265 |
| K01874 | MARS, metG | methionyl-tRNA synthetase [EC:6.1.1.10] | 919 | 162 | 512 | 644 | 560 | 829 | 665 | 661 | 0 | 0 | 0 | 0 | 0 | 413 | 1359 | 0 | 0.002980685 | -7.474950159 |
| K03664 | smpB | SsrA-binding protein | 941 | 149 | 0 | 0 | 518 | 668 | 755 | 428 | 0 | 0 | 0 | 0 | 0 | 0 | 0 | 0 | 0.000250045 | -7.464905347 |
| K07164 | K07164 | uncharacterized protein | 1012 | 784 | 376 | 0 | 260 | 420 | 456 | 0 | 0 | 0 | 0 | 0 | 0 | 0 | 0 | 0 | 0.000226442 | -7.452092857 |
| K00170 | porB | pyruvate ferredoxin oxidoreductase beta subunit [EC:1.2.7.1] | 4897 | 4019 | 2513 | 5641 | 5298 | 5186 | 4713 | 3958 | 0 | 541 | 0 | 0 | 1110 | 1957 | 2076 | 0 | 0.005347711 | -7.450821348 |
| K01999 | livK | branched-chain amino acid transport system substrate-binding | 2516 | 4221 | 3015 | 1913 | 5644 | 4970 | 5644 | 5680 | 0 | 0 | 592 | 832 | 2095 | 924 | 0 | 0 | 0.004597202 | -7.448701194 |
| K11689 | dctQ | C4-dicarboxylate transporter, DctQ subunit | 378 | 180 | 0 | 0 | 706 | 707 | 458 | 831 | 0 | 0 | 0 | 0 | 0 | 0 | 0 | 0 | 0.000250045 | -7.43077614 |
| K06969 | rlmI | 23S rRNA (cytosine1962-C5)-methyltransferase [EC:2.1.1.191] | 486 | 455 | 0 | 0 | 392 | 466 | 455 | 632 | 0 | 0 | 0 | 0 | 0 | 0 | 0 | 0 | 0.000242826 | -7.411346222 |
| K00179 | iorA | indolepyruvate ferredoxin oxidoreductase, alpha subunit [EC:1. | 288 | 297 | 528 | 246 | 338 | 592 | 531 | 629 | 0 | 0 | 0 | 0 | 259 | 308 | 0 | 0 | 0.002044762 | -7.391086192 |
| K03110 | ftsY | fused signal recognition particle receptor | 344 | 396 | 851 | 0 | 250 | 492 | 533 | 302 | 0 | 0 | 0 | 0 | 282 | 0 | 0 | 0 | 0.002429069 | -7.377540838 |
| K03686 | dnaJ | molecular chaperone DnaJ | 0 | 190 | 274 | 0 | 805 | 640 | 622 | 574 | 0 | 0 | 0 | 0 | 0 | 0 | 0 | 0 | 0.000252877 | -7.377178129 |
| K00059 | fabG | 3-oxoacyl-[acyl-carrier protein] reductase [EC:1.1.1.100] | 357 | 778 | 327 | 612 | 646 | 556 | 774 | 558 | 0 | 0 | 0 | 0 | 472 | 0 | 0 | 1758 | 0.003500785 | -7.369376161 |
| K00684 | aat | leucyl/phenylalanyl-tRNA--protein transferase [EC:2.3.2.6] | 0 | 191 | 275 | 514 | 741 | 878 | 594 | 132 | 0 | 0 | 0 | 0 | 182 | 0 | 0 | 0 | 0.002308236 | -7.363625962 |
| K00525 | E1.17.4.1A, nrdA, nrdE | ribonucleoside-diphosphate reductase alpha chain [EC:1.17.4.1 | 879 | 273 | 254 | 147 | 983 | 958 | 1255 | 506 | 0 | 0 | 0 | 0 | 464 | 1452 | 0 | 0 | 0.00340347 | -7.361632525 |
| K10716 | kch, trkA, mthK, pch | voltage-gated potassium channel | 829 | 345 | 253 | 0 | 482 | 632 | 754 | 243 | 0 | 529 | 0 | 0 | 0 | 0 | 0 | 0 | 0.003111253 | -7.357935143 |
| K02669 | pilT | twitching motility protein PilT | 929 | 256 | 0 | 795 | 725 | 710 | 856 | 272 | 0 | 0 | 0 | 0 | 0 | 0 | 2159 | 0 | 0.006969905 | -7.349905252 |
| K01649 | leuA | 2-isopropylmalate synthase [EC:2.3.3.13] | 919 | 2167 | 1003 | 621 | 3474 | 3141 | 2483 | 3024 | 0 | 0 | 0 | 0 | 1344 | 2194 | 0 | 2202 | 0.008143053 | -7.332917599 |
| K08364 | merP | periplasmic mercuric ion binding protein | 769 | 366 | 0 | 1317 | 226 | 311 | 291 | 0 | 0 | 0 | 0 | 0 | 0 | 0 | 0 | 0 | 0.000242826 | -7.330069039 |
| K03531 | ftsZ | cell division protein FtsZ | 1938 | 1143 | 1420 | 4310 | 635 | 577 | 652 | 681 | 0 | 414 | 0 | 1193 | 352 | 0 | 0 | 0 | 0.00368929 | -7.307115576 |
| K03595 | era | GTPase | 0 | 154 | 0 | 416 | 624 | 666 | 565 | 427 | 0 | 0 | 0 | 0 | 0 | 0 | 0 | 0 | 0.000260604 | -7.305952394 |
| K00343 | nuoN | NADH-quinone oxidoreductase subunit N [EC:1.6.5.3] | 0 | 449 | 223 | 580 | 3124 | 2275 | 2214 | 2425 | 0 | 0 | 217 | 0 | 0 | 1086 | 0 | 0 | 0.009387288 | -7.264198718 |
| K00342 | nuoM | NADH-quinone oxidoreductase subunit M [EC:1.6.5.3] | 827 | 558 | 568 | 354 | 2793 | 2506 | 1691 | 790 | 0 | 334 | 0 | 0 | 657 | 442 | 0 | 0 | 0.00353975 | -7.247425128 |
| K02387 | flgB | flagellar basal-body rod protein FlgB | 0 | 155 | 0 | 805 | 496 | 387 | 157 | 828 | 0 | 0 | 0 | 0 | 0 | 0 | 0 | 0 | 0.000296184 | -7.177313708 |
| K00053 | ilvC | ketol-acid reductoisomerase [EC:1.1.1.86] | 441 | 561 | 809 | 0 | 1858 | 1396 | 1821 | 1261 | 0 | 0 | 0 | 0 | 536 | 472 | 0 | 0 | 0.00944117 | -7.138219199 |
| K03272 | gmhC, hldE, waaE, rfaE | D-beta-D-heptose 7-phosphate kinase / D-beta-D-heptose 1-pł | 563 | 153 | 858 | 534 | 659 | 453 | 414 | 0 | 0 | 0 | 0 | 2157 | 0 | 0 | 0 | 0 | 0.007767241 | -7.116311006 |
| K04042 | glmU | bifunctional UDP-N-acetylglucosamine pyrophosphorylase / Gl | 681 | 144 | 624 | 1166 | 152 | 140 | 117 | 586 | 0 | 0 | 0 | 0 | 128 | 454 | 0 | 0 | 0.002481467 | -7.095688521 |
| K03118 | tatC | sec-independent protein translocase protein TatC | 581 | 231 | 532 | 0 | 457 | 324 | 440 | 511 | 0 | 0 | 0 | 0 | 1413 | 0 | 0 | 0 | 0.006255741 | -7.092671816 |
| K01755 | argH, ASL | argininosuccinate lyase [EC:4.3.2.1] | 0 | 150 | 143 | 534 | 346 | 323 | 558 | 210 | 0 | 0 | 0 | 0 | 94 | 0 | 0 | 0 | 0.002110314 | -7.080734484 |
| K01890 | FARSB, pheT | phenylalanyl-tRNA synthetase beta chain [EC:6.1.1.20] | 367 | 373 | 363 | 339 | 487 | 691 | 583 | 87 | 0 | 0 | 0 | 0 | 246 | 785 | 0 | 0 | 0.003106324 | -7.06287207 |
| K02492 | hemA | glutamyl-tRNA reductase [EC:1.2.1.70] | 0 | 273 | 230 | 0 | 148 | 553 | 537 | 535 | 0 | 0 | 0 | 0 | 0 | 0 | 0 | 0 | 0.000296184 | -7.043786286 |
| K03749 | dedD | DedD protein | 0 | 393 | 0 | 530 | 273 | 214 | 257 | 419 | 0 | 0 | 0 | 0 | 0 | 0 | 0 | 0 | 0.000296184 | -7.021782442 |
| K00674 | dapD | 2,3,4,5-tetrahydropyridine-2-carboxylate N-succinyltransferase | 0 | 302 | 618 | 0 | 1567 | 1475 | 1118 | 1587 | 0 | 0 | 0 | 0 | 336 | 0 | 0 | 0 | 0.008669034 | -7.017033999 |
| K00800 | aroA | 3-phosphoshikimate 1-carboxyvinyltransferase [EC:2.5.1.19] | 550 | 349 | 358 | 0 | 398 | 194 | 208 | 0 | 0 | 0 | 0 | 0 | 0 | 0 | 0 | 0 | 0.000267165 | -6.98074944 |
| K02523 | ispB | octaprenyl-diphosphate synthase [EC:2.5.1.90] | 363 | 115 | 0 | 0 | 457 | 408 | 807 | 159 | 0 | 0 | 0 | 0 | 0 | 0 | 0 | 0 | 0.000320522 | -6.973451713 |
| K02409 | fliF | flagellar M-ring protein FliF | 426 | 48 | 116 | 0 | 524 | 669 | 609 | 1181 | 0 | 0 | 0 | 0 | 697 | 0 | 0 | 0 | 0.005843759 | -6.970354503 |
| K09812 | ftsE | cell division transport system ATP-binding protein | 0 | 492 | 0 | 549 | 410 | 244 | 247 | 140 | 0 | 0 | 0 | 0 | 0 | 0 | 0 | 0 | 0.000299891 | -6.960067941 |
| K00077 | panE, apbA | 2-dehydropantoate 2-reductase [EC:1.1.1.169] | 0 | 352 | 485 | 453 | 229 | 328 | 209 | 387 | 0 | 0 | 0 | 0 | 0 | 556 | 0 | 0 | 0.004601339 | -6.958012342 |
| K06881 | nrnA | bifunctional oligoribonuclease and PAP phosphatase NrnA [EC: | 0 | 218 | 420 | 0 | 341 | 308 | 386 | 229 | 0 | 0 | 0 | 0 | 0 | 0 | 0 | 0 | 0.000296184 | -6.943667042 |

| | | | | | | | | | | | | | | | | | | | | |
|---|---|---|---|---|---|---|---|---|---|---|---|---|---|---|---|---|---|---|---|---|
| K03799 | htpX | heat shock protein HtpX [EC:3.4.24.-] | 0 | 113 | 195 | 0 | 756 | 383 | 632 | 169 | 0 | 0 | 0 | 0 | 0 | 0 | 0 | 0 | 0.000339902 | -6.905076477 |
| K01997 | livH | branched-chain amino acid transport system permease proteir | 477 | 265 | 161 | 541 | 354 | 341 | 368 | 209 | 0 | 0 | 0 | 0 | 183 | 1020 | 0 | 0 | 0.003697499 | -6.880956956 |
| K03667 | hslU | ATP-dependent HslUV protease ATP-binding subunit HslU | 0 | 346 | 1598 | 0 | 506 | 871 | 954 | 757 | 0 | 0 | 183 | 0 | 0 | 0 | 0 | 0 | 0.006972324 | -6.873905513 |
| K08223 | fsr | MFS transporter, FSR family, fosmidomycin resistance protein | 0 | 86 | 955 | 0 | 286 | 448 | 319 | 154 | 0 | 0 | 0 | 0 | 0 | 0 | 0 | 0 | 0.000339902 | -6.854710516 |
| K03701 | uvrA | excinuclease ABC subunit A | 120 | 38 | 110 | 0 | 170 | 130 | 145 | 154 | 0 | 0 | 0 | 0 | 0 | 0 | 0 | 0 | 5.5633E-05 | -6.8495216 |
| K03831 | mogA | molybdopterin adenylyltransferase [EC:2.7.7.75] | 0 | 66 | 385 | 720 | 190 | 238 | 264 | 483 | 0 | 0 | 0 | 0 | 255 | 0 | 0 | 0 | 0.003623934 | -6.842445125 |
| K03307 | TC.SSS | solute:Na+ symporter, SSS family | 177 | 141 | 487 | 0 | 469 | 347 | 402 | 295 | 0 | 0 | 0 | 0 | 731 | 0 | 0 | 0 | 0.006268318 | -6.795242597 |
| K03744 | lemA | LemA protein | 0 | 180 | 0 | 1296 | 226 | 205 | 172 | 166 | 0 | 0 | 0 | 0 | 0 | 0 | 0 | 0 | 0.000342307 | -6.761813946 |
| K03741 | ARSC2, arsC | arsenate reductase [EC:1.20.4.1] | 0 | 86 | 1489 | 0 | 225 | 128 | 229 | 266 | 0 | 0 | 0 | 0 | 0 | 0 | 0 | 0 | 0.000362989 | -6.705580754 |
| K03564 | BCP, PRXQ, DOT5 | peroxiredoxin Q/BCP [EC:1.11.1.15] | 0 | 373 | 431 | 805 | 92 | 126 | 142 | 0 | 0 | 0 | 0 | 0 | 0 | 0 | 0 | 0 | 0.000339902 | -6.683322414 |
| K07080 | K07080 | uncharacterized protein | 2251 | 2395 | 2267 | 5007 | 1159 | 1226 | 1330 | 1383 | 0 | 0 | 200 | 0 | 819 | 963 | 0 | 1343 | 0.009477689 | -6.667042649 |
| K02338 | DPO3B, dnaN | DNA polymerase III subunit beta [EC:2.7.7.7] | 216 | 171 | 0 | 0 | 275 | 357 | 245 | 189 | 0 | 0 | 0 | 0 | 0 | 0 | 0 | 0 | 0.000340062 | -6.653761457 |
| K07710 | atoS | two-component system, NtrC family, sensor histidine kinase A | 1060 | 336 | 485 | 0 | 52 | 125 | 128 | 0 | 0 | 0 | 0 | 0 | 0 | 0 | 0 | 0 | 0.000362989 | -6.612922716 |
| K01719 | hemD, UROS | uroporphyrinogen-III synthase [EC:4.2.1.75] | 0 | 229 | 331 | 0 | 283 | 229 | 164 | 158 | 0 | 0 | 0 | 0 | 0 | 0 | 0 | 0 | 0.000342307 | -6.602580457 |
| K01358 | clpP, CLPP | ATP-dependent Clp protease, protease subunit [EC:3.4.21.92] | 0 | 106 | 0 | 0 | 827 | 972 | 1832 | 1436 | 0 | 0 | 0 | 0 | 0 | 0 | 0 | 0 | 0.001969265 | -6.586848455 |
| K03978 | engB | GTP-binding protein | 0 | 705 | 1000 | 0 | 462 | 582 | 633 | 388 | 0 | 0 | 0 | 0 | 234 | 0 | 0 | 0 | 0.009899258 | -6.584293723 |
| K00573 | E2.1.1.77, pcm | protein-L-isoaspartate(D-aspartate) O-methyltransferase [EC:2 | 0 | 105 | 0 | 586 | 130 | 219 | 185 | 291 | 0 | 0 | 0 | 0 | 0 | 0 | 0 | 0 | 0.000362989 | -6.550358838 |
| K01657 | trpE | anthranilate synthase component I [EC:4.1.3.27] | 241 | 148 | 0 | 0 | 91 | 230 | 298 | 395 | 0 | 0 | 0 | 0 | 0 | 0 | 0 | 0 | 0.000362989 | -6.538547422 |
| K03641 | tolB | TolB protein | 0 | 87 | 178 | 0 | 250 | 354 | 339 | 171 | 0 | 0 | 0 | 0 | 0 | 0 | 0 | 0 | 0.000362989 | -6.519333731 |
| K00014 | aroE | shikimate dehydrogenase [EC:1.1.1.25] | 279 | 88 | 0 | 0 | 190 | 107 | 272 | 366 | 0 | 0 | 0 | 0 | 0 | 0 | 0 | 0 | 0.000392953 | -6.436254635 |
| K01079 | serB, PSPH | phosphoserine phosphatase [EC:3.1.3.3] | 611 | 237 | 0 | 0 | 102 | 151 | 100 | 194 | 0 | 0 | 0 | 0 | 0 | 0 | 0 | 0 | 0.000397391 | -6.40923555 |
| K01993 | ABC-2.TX | HlyD family secretion protein | 715 | 162 | 468 | 0 | 50 | 123 | 108 | 0 | 0 | 0 | 0 | 0 | 0 | 0 | 0 | 0 | 0.00040727 | -6.3643187 |
| K01462 | PDF, def | peptide deformylase [EC:3.5.1.88] | 0 | 204 | 0 | 0 | 798 | 782 | 768 | 535 | 0 | 0 | 0 | 0 | 0 | 0 | 0 | 0 | 0.001969265 | -6.322407021 |
| K11473 | glcF | glycolate oxidase iron-sulfur subunit | 0 | 80 | 0 | 0 | 1219 | 779 | 646 | 976 | 0 | 0 | 0 | 0 | 0 | 0 | 0 | 0 | 0.002057842 | -6.308486918 |
| K11784 | mqnC | cyclic dehypoxanthinyl futalosine synthase [EC:1.21.98.1] | 205 | 65 | 0 | 0 | 257 | 178 | 218 | 180 | 0 | 0 | 0 | 0 | 0 | 0 | 0 | 0 | 0.000412148 | -6.30417102 |
| K03072 | secD | preprotein translocase subunit SecD | 0 | 153 | 264 | 0 | 763 | 700 | 744 | 195 | 0 | 0 | 0 | 0 | 108 | 0 | 0 | 0 | 0.009410625 | -6.238838871 |
| K03570 | mreC | rod shape-determining protein MreC | 441 | 210 | 404 | 0 | 43 | 102 | 100 | 0 | 0 | 0 | 0 | 0 | 0 | 0 | 0 | 0 | 0.000426416 | -6.223441022 |
| K01972 | E6.5.1.2, ligA, ligB | DNA ligase (NAD+) [EC:6.5.1.2] | 0 | 198 | 0 | 0 | 937 | 728 | 625 | 325 | 0 | 0 | 0 | 0 | 0 | 0 | 0 | 0 | 0.002034768 | -6.205115998 |
| K00339 | nuoJ | NADH-quinone oxidoreductase subunit J [EC:1.6.5.3] | 0 | 161 | 0 | 0 | 993 | 978 | 806 | 201 | 0 | 0 | 0 | 0 | 0 | 0 | 0 | 0 | 0.002057842 | -6.190475405 |
| K03074 | secF | preprotein translocase subunit SecF | 0 | 167 | 0 | 0 | 1034 | 641 | 780 | 231 | 0 | 0 | 0 | 0 | 0 | 0 | 0 | 0 | 0.002057842 | -6.147265205 |
| K00104 | glcD | glycolate oxidase [EC:1.1.3.15] | 0 | 469 | 0 | 0 | 323 | 396 | 418 | 785 | 0 | 0 | 0 | 0 | 0 | 0 | 0 | 0 | 0.002034768 | -6.145222076 |
| K00788 | thiE | thiamine-phosphate pyrophosphorylase [EC:2.5.1.3] | 0 | 467 | 0 | 0 | 345 | 656 | 661 | 278 | 0 | 0 | 0 | 0 | 0 | 0 | 0 | 0 | 0.002034768 | -6.141680031 |
| K02412 | fliI | flagellum-specific ATP synthase [EC:3.6.3.14] | 0 | 66 | 0 | 0 | 766 | 777 | 796 | 592 | 0 | 0 | 0 | 0 | 0 | 0 | 0 | 0 | 0.002163997 | -6.136061765 |
| K04486 | E3.1.3.15B | histidinol-phosphatase (PHP family) [EC:3.1.3.15] | 277 | 88 | 0 | 0 | 86 | 65 | 194 | 255 | 0 | 0 | 0 | 0 | 0 | 0 | 0 | 0 | 0.00050084 | -6.076016332 |
| K02337 | DPO3A1, dnaE | DNA polymerase III subunit alpha [EC:2.7.7.7] | 0 | 72 | 111 | 0 | 306 | 181 | 267 | 53 | 0 | 0 | 0 | 0 | 0 | 0 | 0 | 0 | 0.000507677 | -6.061452466 |
| K01935 | bioD | dethiobiotin synthetase [EC:6.3.3.3] | 0 | 63 | 0 | 680 | 117 | 62 | 116 | 151 | 0 | 0 | 0 | 0 | 0 | 0 | 0 | 0 | 0.000550407 | -6.036740016 |
| K01599 | hemE, UROD | uroporphyrinogen decarboxylase [EC:4.1.1.37] | 0 | 94 | 0 | 0 | 782 | 678 | 507 | 330 | 0 | 0 | 0 | 0 | 0 | 0 | 0 | 0 | 0.002175467 | -5.990339912 |
| K01653 | E2.2.1.6S, ilvH, ilvN | acetolactate synthase I/III small subunit [EC:2.2.1.6] | 0 | 0 | 618 | 0 | 509 | 299 | 392 | 227 | 0 | 0 | 0 | 0 | 0 | 0 | 0 | 0 | 0.002069187 | -5.988461644 |
| K03182 | ubiD | 4-hydroxy-3-polyprenylbenzoate decarboxylase [EC:4.1.1.98] | 0 | 171 | 990 | 0 | 333 | 529 | 296 | 0 | 0 | 0 | 0 | 0 | 0 | 0 | 0 | 0 | 0.002049571 | -5.987219621 |
| K06920 | queC | 7-cyano-7-deazaguanine synthase [EC:6.3.4.20] | 0 | 0 | 508 | 0 | 272 | 226 | 321 | 689 | 0 | 0 | 0 | 0 | 0 | 0 | 0 | 0 | 0.002120289 | -5.955145553 |
| K01270 | pepD | dipeptidase D [EC:3.4.13.-] | 350 | 0 | 0 | 600 | 318 | 278 | 281 | 0 | 0 | 0 | 0 | 0 | 0 | 0 | 0 | 0 | 0.002049571 | -5.893026549 |
| K01892 | HARS, hisS | histidyl-tRNA synthetase [EC:6.1.1.21] | 0 | 337 | 0 | 0 | 455 | 279 | 319 | 299 | 0 | 0 | 0 | 0 | 0 | 0 | 0 | 0 | 0.002163997 | -5.859900924 |
| K01693 | hisB | imidazoleglycerol-phosphate dehydratase [EC:4.2.1.19] | 0 | 239 | 0 | 0 | 254 | 279 | 448 | 496 | 0 | 0 | 0 | 0 | 0 | 0 | 0 | 0 | 0.002175467 | -5.846139436 |
| K01151 | nfo | deoxyribonuclease IV [EC:3.1.21.2] | 0 | 94 | 0 | 1020 | 359 | 328 | 311 | 0 | 0 | 0 | 0 | 0 | 0 | 0 | 0 | 0 | 0.002175467 | -5.822337027 |
| K00231 | PPOX, hemY | oxygen-dependent protoporphyrinogen oxidase [EC:1.3.3.4] | 0 | 245 | 0 | 0 | 474 | 440 | 270 | 231 | 0 | 0 | 0 | 0 | 0 | 0 | 0 | 0 | 0.002186808 | -5.815248982 |
| K07095 | K07095 | uncharacterized protein | 511 | 188 | 0 | 0 | 307 | 347 | 330 | 0 | 0 | 0 | 0 | 0 | 0 | 0 | 0 | 0 | 0.002091502 | -5.81508711 |
| K04744 | lptD, imp, ostA | LPS-assembly protein | 0 | 0 | 0 | 376 | 300 | 192 | 305 | 349 | 0 | 0 | 0 | 0 | 0 | 0 | 0 | 0 | 0.002253234 | -5.757139426 |
| K01703 | leuC | 3-isopropylmalate/(R)-2-methylmalate dehydratase large subu | 0 | 27 | 0 | 0 | 723 | 565 | 721 | 213 | 0 | 0 | 0 | 0 | 0 | 0 | 0 | 0 | 0.00251659 | -5.705127948 |
| K02078 | acpP | acyl carrier protein | 0 | 437 | 840 | 0 | 180 | 176 | 138 | 0 | 0 | 0 | 0 | 0 | 0 | 0 | 0 | 0 | 0.002264575 | -5.679639143 |
| K03304 | tehA | tellurite resistance protein | 426 | 0 | 0 | 0 | 286 | 108 | 163 | 660 | 0 | 0 | 0 | 0 | 0 | 0 | 0 | 0 | 0.002331252 | -5.670223294 |
| K03543 | emrA | membrane fusion protein, multidrug efflux system | 0 | 116 | 0 | 0 | 166 | 356 | 628 | 328 | 0 | 0 | 0 | 0 | 0 | 0 | 0 | 0 | 0.002331252 | -5.669591771 |
| K00991 | ispD | 2-C-methyl-D-erythritol 4-phosphate cytidylyltransferase [EC:2. | 294 | 131 | 0 | 0 | 28 | 78 | 62 | 129 | 0 | 0 | 0 | 0 | 0 | 0 | 0 | 0 | 0.000766079 | -5.661296503 |
| K07461 | K07461 | putative endonuclease | 0 | 0 | 0 | 0 | 1113 | 856 | 1782 | 1533 | 0 | 0 | 0 | 0 | 0 | 0 | 0 | 0 | 0.007071436 | -5.658366781 |
| K00790 | murA | UDP-N-acetylglucosamine 1-carboxyvinyltransferase [EC:2.5.1. | 0 | 220 | 0 | 0 | 385 | 263 | 311 | 152 | 0 | 0 | 0 | 0 | 0 | 0 | 0 | 0 | 0.002331252 | -5.615250414 |
| K00930 | argB | acetylglutamate kinase [EC:2.7.2.8] | 542 | 86 | 0 | 0 | 292 | 220 | 348 | 0 | 0 | 0 | 0 | 0 | 0 | 0 | 0 | 0 | 0.002331252 | -5.603998692 |
| K01042 | selA | L-seryl-tRNA(Ser) seleniumtransferase [EC:2.9.1.1] | 0 | 372 | 744 | 0 | 159 | 199 | 93 | 0 | 0 | 0 | 0 | 0 | 0 | 0 | 0 | 0 | 0.002331252 | -5.557641394 |
| K00872 | thrB1 | homoserine kinase [EC:2.7.1.39] | 0 | 102 | 0 | 0 | 387 | 178 | 481 | 206 | 0 | 0 | 0 | 0 | 0 | 0 | 0 | 0 | 0.002394914 | -5.541792072 |
| K02527 | kdtA, waaA | 3-deoxy-D-manno-octulosonic-acid transferase [EC:2.4.99.12 2 | 0 | 120 | 0 | 648 | 237 | 190 | 201 | 0 | 0 | 0 | 0 | 0 | 0 | 0 | 0 | 0 | 0.002331252 | -5.532542321 |
| K02372 | fabZ | 3-hydroxyacyl-[acyl-carrier-protein] dehydratase [EC:4.2.1.59] | 0 | 191 | 0 | 0 | 177 | 323 | 227 | 264 | 0 | 0 | 0 | 0 | 0 | 0 | 0 | 0 | 0.002331252 | -5.530128912 |
| K04485 | radA, sms | DNA repair protein RadA/Sms | 0 | 227 | 0 | 0 | 258 | 301 | 370 | 100 | 0 | 0 | 0 | 0 | 0 | 0 | 0 | 0 | 0.002357637 | -5.528616664 |
| K02415 | fliL | flagellar FliL protein | 0 | 194 | 0 | 0 | 160 | 278 | 277 | 268 | 0 | 0 | 0 | 0 | 0 | 0 | 0 | 0 | 0.002331252 | -5.526322577 |
| K07173 | luxS | S-ribosylhomocysteine lyase [EC:4.4.1.21] | 0 | 258 | 372 | 0 | 394 | 78 | 228 | 0 | 0 | 0 | 0 | 0 | 0 | 0 | 0 | 0 | 0.002331252 | -5.523895105 |
| K11754 | folC | dihydrofolate synthase / folylpolyglutamate synthase [EC:6.3.2 | 0 | 283 | 504 | 0 | 221 | 217 | 83 | 0 | 0 | 0 | 0 | 0 | 0 | 0 | 0 | 0 | 0.002331252 | -5.492917851 |
| K01012 | bioB | biotin synthase [EC:2.8.1.6] | 260 | 196 | 0 | 0 | 285 | 261 | 134 | 0 | 0 | 0 | 0 | 0 | 0 | 0 | 0 | 0 | 0.002331252 | -5.473868076 |
| K02115 | ATPF1G, atpG | F-type H+-transporting ATPase subunit gamma | 0 | 164 | 0 | 0 | 217 | 378 | 164 | 214 | 0 | 0 | 0 | 0 | 0 | 0 | 0 | 0 | 0.002394914 | -5.471246672 |
| K01775 | alr | alanine racemase [EC:5.1.1.1] | 0 | 238 | 0 | 376 | 115 | 262 | 175 | 0 | 0 | 0 | 0 | 0 | 0 | 0 | 0 | 0 | 0.002331252 | -5.460142178 |
| K03813 | modD | molybdenum transport protein [EC:2.4.2.-] | 374 | 368 | 0 | 0 | 143 | 240 | 97 | 0 | 0 | 0 | 0 | 0 | 0 | 0 | 0 | 0 | 0.002331252 | -5.45481535 |
| K00794 | ribH, RIB4 | 6,7-dimethyl-8-ribityllumazine synthase [EC:2.5.1.78] | 500 | 0 | 0 | 0 | 109 | 185 | 191 | 222 | 0 | 0 | 0 | 0 | 0 | 0 | 0 | 0 | 0.002429069 | -5.45321356 |
| K03833 | selB, EEFSEC | selenocysteine-specific elongation factor | 0 | 220 | 624 | 0 | 163 | 120 | 169 | 0 | 0 | 0 | 0 | 0 | 0 | 0 | 0 | 0 | 0.002331252 | -5.452433603 |
| K15738 | uup | ATP-binding cassette, subfamily F, uup | 0 | 196 | 0 | 0 | 310 | 242 | 127 | 218 | 0 | 0 | 0 | 0 | 0 | 0 | 0 | 0 | 0.002404388 | -5.444555674 |
| K01585 | speA | arginine decarboxylase [EC:4.1.1.19] | 0 | 25 | 0 | 0 | 241 | 455 | 320 | 438 | 0 | 0 | 0 | 0 | 0 | 0 | 0 | 0 | 0.002708317 | -5.438684044 |
| K02341 | DPO3D2, holB | DNA polymerase III subunit delta' [EC:2.7.7.7] | 0 | 232 | 0 | 0 | 222 | 223 | 195 | 170 | 0 | 0 | 0 | 0 | 0 | 0 | 0 | 0 | 0.002400971 | -5.432109698 |
| K09141 | K09141 | uncharacterized protein | 0 | 166 | 0 | 0 | 180 | 236 | 324 | 163 | 0 | 0 | 0 | 0 | 0 | 0 | 0 | 0 | 0.002404388 | -5.428400564 |
| K06966 | K06966 | uncharacterized protein | 0 | 0 | 567 | 0 | 230 | 167 | 93 | 136 | 0 | 0 | 0 | 0 | 0 | 0 | 0 | 0 | 0.002504319 | -5.373141604 |
| K18929 | lldF | L-lactate dehydrogenase complex protein LldF | 0 | 0 | 296 | 0 | 229 | 125 | 113 | 285 | 0 | 0 | 0 | 0 | 0 | 0 | 0 | 0 | 0.002481467 | -5.372635743 |
| K07391 | comM | magnesium chelatase family protein | 0 | 96 | 0 | 0 | 106 | 232 | 274 | 359 | 0 | 0 | 0 | 0 | 0 | 0 | 0 | 0 | 0.002546909 | -5.345310822 |
| K07066 | K07066 | uncharacterized protein | 554 | 176 | 0 | 0 | 217 | 138 | 83 | 0 | 0 | 0 | 0 | 0 | 0 | 0 | 0 | 0 | 0.002429795 | -5.340267205 |
| K07114 | yfbK | Ca-activated chloride channel homolog | 0 | 46 | 0 | 0 | 425 | 284 | 227 | 176 | 0 | 0 | 0 | 0 | 0 | 0 | 0 | 0 | 0.002613702 | -5.337327073 |
| K00566 | mnmA, trmU, TRMU | tRNA-specific 2-thiouridylase [EC:2.8.1.-] | 0 | 93 | 0 | 0 | 134 | 294 | 246 | 224 | 0 | 0 | 0 | 0 | 0 | 0 | 0 | 0 | 0.002544448 | -5.319568624 |
| K02016 | ABC.FEV.S | iron complex transport system substrate-binding protein | 0 | 38 | 0 | 0 | 150 | 246 | 336 | 390 | 0 | 0 | 0 | 0 | 0 | 0 | 0 | 0 | 0.002726755 | -5.30510563 |
| K03642 | rlpA | rare lipoprotein A | 0 | 117 | 0 | 0 | 115 | 151 | 539 | 153 | 0 | 0 | 0 | 0 | 0 | 0 | 0 | 0 | 0.002611491 | -5.285753278 |
| K01791 | wecB | UDP-N-acetylglucosamine 2-epimerase (non-hydrolysing) [EC:5 | 0 | 0 | 0 | 335 | 135 | 119 | 99 | 305 | 0 | 0 | 0 | 0 | 0 | 0 | 0 | 0 | 0.002611491 | -5.280432726 |
| K12410 | npdA | NAD-dependent deacetylase [EC:3.5.1.-] | 0 | 74 | 0 | 0 | 128 | 204 | 279 | 275 | 0 | 0 | 0 | 0 | 0 | 0 | 0 | 0 | 0.002613702 | -5.264970599 |
| K07665 | cusR, copR, silR | two-component system, OmpR family, copper resistance phos | 0 | 0 | 0 | 0 | 768 | 777 | 1029 | 361 | 0 | 0 | 0 | 0 | 0 | 0 | 0 | 0 | 0.008616462 | -5.214653094 |
| K09817 | znuC | zinc transport system ATP-binding protein [EC:3.6.3.-] | 0 | 113 | 326 | 0 | 0 | 170 | 130 | 130 | 0 | 0 | 0 | 0 | 0 | 0 | 0 | 0 | 0.002595405 | -5.198073707 |
| K15497 | wtpC | molybdate/tungstate transport system ATP-binding protein [E | 0 | 140 | 0 | 0 | 218 | 235 | 66 | 194 | 0 | 0 | 0 | 0 | 0 | 0 | 0 | 0 | 0.002696211 | -5.178530064 |
| K18429 | legG, neuC2 | GDP/UDP-N,N'-diacetylbacillosamine 2-epimerase (hydrolysin | 299 | 0 | 274 | 0 | 226 | 76 | 66 | 0 | 0 | 0 | 0 | 0 | 0 | 0 | 0 | 0 | 0.002611491 | -5.167028097 |
| K09816 | znuB | zinc transport system permease protein | 0 | 109 | 0 | 0 | 119 | 150 | 173 | 234 | 0 | 0 | 0 | 0 | 0 | 0 | 0 | 0 | 0.002696211 | -5.15145213 |
| K07456 | mutS2 | DNA mismatch repair protein MutS2 | 0 | 236 | 407 | 0 | 93 | 123 | 71 | 0 | 0 | 0 | 0 | 0 | 0 | 0 | 0 | 0 | 0.002613702 | -5.134931598 |
| K06193 | phnA | protein PhnA | 404 | 64 | 0 | 0 | 0 | 93 | 152 | 177 | 0 | 0 | 0 | 0 | 0 | 0 | 0 | 0 | 0.002808561 | -5.111953368 |

| | | | | | | | | | | | | | | | | | | | | |
|---|---|---|---|---|---|---|---|---|---|---|---|---|---|---|---|---|---|---|---|---|
| K08679 | E5.1.3.6 | UDP-glucuronate 4-epimerase [EC:5.1.3.6] | 0 | 41 | 0 | 1083 | 186 | 85 | 95 | 0 | 0 | 0 | 0 | 0 | 0 | 0 | 0 | 0 | 0.002991887 | -5.108821003 |
| K09774 | lptA | lipopolysaccharide export system protein LptA | 0 | 0 | 528 | 0 | 56 | 79 | 79 | 293 | 0 | 0 | 0 | 0 | 0 | 0 | 0 | 0 | 0.002991887 | -5.084157611 |
| K02428 | rdgB | XTP/dITP diphosphohydrolase [EC:3.6.1.66] | 0 | 57 | 0 | 0 | 106 | 144 | 123 | 397 | 0 | 0 | 0 | 0 | 0 | 0 | 0 | 0 | 0.002991887 | -5.04380412 |
| K01494 | dcd | dCTP deaminase [EC:3.5.4.13] | 0 | 71 | 0 | 0 | 88 | 111 | 291 | 198 | 0 | 0 | 0 | 0 | 0 | 0 | 0 | 0 | 0.002991887 | -5.031501375 |
| K03639 | MOCS1, moaA | cyclic pyranopterin phosphate synthase [EC:4.1.99.18] | 0 | 189 | 0 | 0 | 92 | 187 | 54 | 210 | 0 | 0 | 0 | 0 | 0 | 0 | 0 | 0 | 0.002991887 | -5.016400072 |
| K15496 | wtpB | molybdate/tungstate transport system permease protein | 0 | 89 | 0 | 0 | 165 | 119 | 170 | 123 | 0 | 0 | 0 | 0 | 0 | 0 | 0 | 0 | 0.002912306 | -5.014231197 |
| K01992 | ABC-2.P | ABC-2 type transport system permease protein | 0 | 31 | 0 | 340 | 114 | 164 | 176 | 0 | 0 | 0 | 0 | 0 | 0 | 0 | 0 | 0 | 0.002991887 | -4.991820068 |
| K08306 | mltC | membrane-bound lytic murein transglycosylase C | 0 | 0 | 0 | 0 | 367 | 398 | 565 | 748 | 0 | 0 | 0 | 0 | 0 | 0 | 0 | 0 | 0.009477689 | -4.989457353 |
| K02169 | bioC | malonyl-CoA O-methyltransferase [EC:2.1.1.197] | 0 | 46 | 0 | 0 | 117 | 135 | 158 | 259 | 0 | 0 | 0 | 0 | 0 | 0 | 0 | 0 | 0.003022376 | -4.980135272 |
| K13292 | lgt, umpA | phosphatidylglycerol:prolipoprotein diacylglycerol transferase [ | 284 | 0 | 0 | 0 | 141 | 86 | 64 | 135 | 0 | 0 | 0 | 0 | 0 | 0 | 0 | 0 | 0.002991887 | -4.977479546 |
| K00833 | bioA | adenosylmethionine---8-amino-7-oxononanoate aminotransfe | 0 | 95 | 0 | 0 | 86 | 163 | 134 | 123 | 0 | 0 | 0 | 0 | 0 | 0 | 0 | 0 | 0.002991887 | -4.923561218 |
| K02238 | comEC | competence protein ComEC | 0 | 56 | 0 | 0 | 192 | 127 | 100 | 154 | 0 | 0 | 0 | 0 | 0 | 0 | 0 | 0 | 0.003039777 | -4.917218033 |
| K00640 | cysE | serine O-acetyltransferase [EC:2.3.1.30] | 0 | 0 | 282 | 0 | 30 | 83 | 210 | 135 | 0 | 0 | 0 | 0 | 0 | 0 | 0 | 0 | 0.003248763 | -4.906405385 |
| K02394 | flgI | flagellar P-ring protein precursor FlgI | 0 | 0 | 0 | 0 | 278 | 444 | 390 | 777 | 0 | 0 | 0 | 0 | 0 | 0 | 0 | 0 | 0.009920763 | -4.901151584 |
| K01740 | E2.5.1.49, metY | O-acetylhomoserine (thiol)-lyase [EC:2.5.1.49] | 0 | 57 | 332 | 0 | 157 | 76 | 82 | 0 | 0 | 0 | 0 | 0 | 0 | 0 | 0 | 0 | 0.002991887 | -4.876667564 |
| K02020 | modA | molybdate transport system substrate-binding protein | 0 | 47 | 0 | 0 | 176 | 133 | 96 | 131 | 0 | 0 | 0 | 0 | 0 | 0 | 0 | 0 | 0.003216853 | -4.843303019 |
| K02013 | ABC.FEV.A | iron complex transport system ATP-binding protein [EC:3.6.3.3 | 0 | 86 | 0 | 0 | 63 | 122 | 41 | 238 | 0 | 0 | 0 | 0 | 0 | 0 | 0 | 0 | 0.003607369 | -4.70859514 |
| K11753 | ribF | riboflavin kinase / FMN adenylyltransferase [EC:2.7.1.26 2.7.7. | 0 | 44 | 0 | 0 | 110 | 128 | 42 | 246 | 0 | 0 | 0 | 0 | 0 | 0 | 0 | 0 | 0.003666263 | -4.708410413 |
| K06153 | bacA | undecaprenyl-diphosphatase [EC:3.6.1.27] | 0 | 125 | 0 | 0 | 38 | 106 | 59 | 173 | 0 | 0 | 0 | 0 | 0 | 0 | 0 | 0 | 0.003639604 | -4.667095992 |
| K03743 | pncC | nicotinamide-nucleotide amidase [EC:3.5.1.42] | 0 | 72 | 0 | 0 | 101 | 97 | 34 | 201 | 0 | 0 | 0 | 0 | 0 | 0 | 0 | 0 | 0.00368929 | -4.656834888 |
| K15778 | pmm-pgm | phosphomannomutase / phosphoglucomutase [EC:5.4.2.8 5.4.2 | 0 | 80 | 0 | 0 | 46 | 57 | 89 | 209 | 0 | 0 | 0 | 0 | 0 | 0 | 0 | 0 | 0.003739095 | -4.619118515 |
| K12287 | mshQ | MSHA biogenesis protein MshQ | 0 | 0 | 276 | 0 | 59 | 40 | 22 | 132 | 0 | 0 | 0 | 0 | 0 | 0 | 0 | 0 | 0.00459647 | -4.489110712 |
| K02703 | psbA | photosystem II P680 reaction center D1 protein | 0 | 0 | 0 | 0 | 0 | 0 | 0 | 0 | 0 | 1801 | 339 | 0 | 231 | 814 | 0 | 4545 | 0.001736484 | 6.748478666 |
